# Insights into Cannabinoid Receptor 2 (CB2) anterograde trafficking and pharmacological chaperoning

**DOI:** 10.1101/2025.02.23.639698

**Authors:** Caitlin RM Oyagawa, Braden Woodhouse, Karren C Wood, Michelle Glass, Natasha L Grimsey

## Abstract

Cannabinoid Receptor 2 (CB_2_) is a promising therapeutic target for modulating inflammation. Canonical signalling responses to receptor ligands are critically dependent on cell surface receptor expression. However, it is also now appreciated that intracellular G protein-coupled receptors can contribute to signalling responses and influence functional outcomes. Therefore, understanding how the subcellular distribution of receptors is controlled is also highly pertinent. CB_2_ is observed to be expressed at the cell surface as well as having a considerable proportion expressed intracellularly. Despite this distribution being well established, little is known about the regulation of CB_2_ anterograde trafficking and subcellular distribution. We report that sustained treatment with a range of CB_2_ agonists and inverse agonists stimulates a distinct population of CB_2_ to be delivered to the cell surface, at various expression levels and despite agonists concurrently internalising cell surface CB_2_. We present evidence that this ligand-stimulated anterograde trafficking is a result of CB_2_ agonists, as well as inverse agonists, acting as pharmacological chaperones. We also report that a di-lysine (KK) motif in the CB_2_ C-terminal tail is required for basal delivery to the cell surface. Corroborating the hypothesis that CB_2_ ligands can act as pharmacological chaperones, sustained CB_2_ ligand stimulation induces cell surface expression of the mutated receptor and alters maturation states as measured by western blotting. Our finding that prolonged exposure to CB_2_ ligands can induce CB_2_ cell surface delivery via pharmacological chaperoning may well have important implications for optimal design of CB_2_-targeted therapeutics.

## 1 Introduction

Cannabinoid Receptor 2 (CB_2_) is a class A G protein-coupled receptor (GPCR) that is predominantly expressed in immune tissues and cells, with functions including modulation of immune cell migration, chemotaxis, differentiation, and synthesis of cytokines, resulting in predominantly immunosuppressive and anti-inflammatory effects [1,2]. As such, CB_2_ has been indicated as a potential therapeutic target in chronic inflammatory conditions including osteoporosis, inflammatory bowel disease, and neurodegenerative diseases involving neuroinflammation [1–3]. Due to its largely peripheral and non-neuronal distribution, activation of CB_2_ is not associated with the psychotropic effects of cannabis, further supporting it as a promising therapeutic target.

GPCRs have traditionally been considered to primarily function at the cell surface, where their activation by extracellular ligands results in the transduction of downstream signals. Following either ligand-induced or constitutive activation, the majority of GPCRs desensitise and are subsequently internalised. CB_2_ canonically couples to Gα_i/o_ G proteins [4] and has been demonstrated to internalise following activation [5–8] though this has been suggested to be dependent upon the specific ligand [7,9]. Post-endocytic trafficking is a critical contributor to the subsequent potential for re-sensitisation to receptor ligands, with CB_2_ understood to be recycled back to the cell surface as opposed to degraded [5,6].

A range of GPCRs have also now been reported to also be expressed intracellularly in unstimulated cells, and activation by cell permeable ligands with initiation of non-canonical signalling has been observed for a number of these, with implications for downstream function [10,11]. This is also true for CB_2_, as a number of studies have found that a considerable proportion of CB_2_ is expressed intracellularly in both immortalised cell lines and human primary immune cells [5,7,12–15]. Two studies to date have reported the potential for signalling via intracellular CB_2_, one indicating Gα_q_ coupling from CB_2_ in endolysosomes [12] and the other inferring intracellular signalling in rat medial prefrontal cortex pyramidal neurons based on predominant intracellular CB_2_ expression and timing differences of signalling onset, depending on whether ligands were applied extracellularly or intracellularly [15].

Intracellular GPCR expression has also been associated with abnormal signalling and trafficking, in particular with relation to receptors that require interaction with an accessory protein in order to be delivered to the cell surface, and naturally occurring receptor mutations that prevent cell surface expression. In the latter case, the interaction of cell-permeable ligands with intracellular receptors can act as “pharmacological chaperones” (also referred to as “pharmacoperones”) to facilitate receptor delivery to the cell surface and enable functional recovery [16,17]. This is hypothesised to be via facilitation of appropriate receptor folding and/or conformation which enables subsequent progression through the secretory pathway [18]. Pharmacological chaperoning of intracellularly retained receptor mutants has been shown for several GPCRs, including the melanocortin-4 receptor [19], µ-opioid receptor [20], and V2 vasopressin receptor [21]. Interestingly, surface expression of wild-type receptors can also be modulated via pharmacological chaperoning [22]. Signalling responses have also been implicated in the modulation of cell surface delivery of proteins, acting as a form of positive feedback with the effect of maintaining ongoing responsiveness [23,24].

Given the potential for differential signalling responses between subcellular receptor populations, regulation of the subcellular distribution of GPCRs is likely to be a critical determinant of functional outcome of GPCR activation. In this study, we undertook to further understand CB_2_ intracellular trafficking and subcellular distribution. Herein we describe our findings that CB_2_ ligands can act as pharmacological chaperones of newly synthesised CB_2_, and that this trafficking pathway is regulated distinctly from basal and post-endocytic subcellular distribution.

## 2 Methods

### 2.1 Drugs and chemicals

CP 55,940, anandamide (AEA), 2-arachidonoyl glycerol (2-AG) were from Cayman Chemical (Ann Arbor, MI, USA). Isoproterenol, salbutamol, pertussis toxin (PTX), U73122, LY294002, GF109203X hydrochloride, monensin and cycloheximide (CHX) were from Sigma-Aldrich (St. Louis, MO). WIN 55,212-2 and HU308 were from Tocris Bioscience (Bristol, UK), and Δ^9^tetrahydrocannabinol (Δ^9^-THC) was from THC Pharm (Frankfurt, Germany). AM630 and SR144528 were kind gifts from Roche (Basel, Switzerland). Gallein was from Santa Cruz Biotechnology (Dallas, TX, USA) and brefeldin A (BFA) from BioLegend (San Diego, CA, USA).

### 2.2 Generation of stable cell lines and cell culture

HEK Flp-in-293 cells (Thermo Fisher Scientific, Waltham, MA; #R750-07) were stably or transiently transfected to express human (h)CB_2_. This hCB_2_ wild-type (wt) construct (hCB_2_ wt), with an haemagglutinin (HA) tag and three thrombin cleavage sites (3TCS) appended at the N-terminus of the receptor, was assembled by overlap-extension PCR, using hCB_2_ sequence from the cDNA Resource Centre (www.cdna.org, CNR0200000; 63Q single nucleotide polymorphism [25]) and synthetic DNA oligos for the tags (Integrated DNA Technologies, Coralville, IA), then ligated into pcDNA5/FRT vector (Thermo Fisher Scientific) via NheI and XhoI restriction sites using standard molecular biology techniques, and sequence verified (see also [26], pERK section). hCB_2_ KK318-9EE (“KK mutant”), ETE339-41ATA (“ETE mutant”) and truncation of the last 12 amino acids from the C-terminus (Δ12) were generated via site-directed mutagenesis utilising the Quikchange^®^ approach (Stratagene, San Diego, CA, USA) with slight modifications and using KAPA HiFi polymerase (KAPA Biosystems, South Africa). The HA-3TCS-hβ_2_-AR pcDNA5/FRT (hβ_2_-AR) plasmid was generated with an analogous strategy as for hCB_2_ wt (www.cdna.org, AR0B20TN00).

Cell lines stably expressing hCB_2_ or hβ_2_-AR plasmids were generated by co-transfecting the receptor-containing plasmid with pOG44 (Thermo Fisher Scientific) to facilitate targeted integration into the FRT site of the HEK Flp-in-293 cell line using Lipofectamine 2000 (Thermo Fisher Scientific). Hygromycin B (Thermo Fisher Scientific) was used for initial selection of transfected cells (100 µg·mL^-1^) and subsequent routine maintenance (50 µg·mL^-1^). The 3HA-hCB_1_ construct and HEK cell line have been described previously [27].

All cell lines were maintained in high-glucose DMEM (HyClone, GE Healthcare Life Sciences, Chicago, IL, USA) supplemented with 10% FBS (NZ-origin; Moregate Biotech, Brisbane, Australia) and appropriate selection antibiotics, and incubated at 37°C in 5% CO_2_ in a humidified atmosphere.

### 2.3 Immunocytochemistry and receptor trafficking assays

Cells were seeded at a density of 45,000 cells/well (for stably expressing lines) or 30,000 cells/well (for cells to be transiently transfected) in poly-D-lysine-treated (PDL; Sigma-Aldrich) clear 96-well plates (Nunc, Thermo Fisher Scientific). For assays involving transient receptor expression, receptor constructs were transfected approximately 24 hours post-seeding using Lipofectamine 2000 (Thermo Fisher Scientific), per a slightly modified manufacturer’s protocol.

Assays were carried out approximately 24 hours post-seeding for stable cell lines and 18-24 hours post-transfection for transiently transfected cells. All drugs and reagents were prepared in high glucose DMEM supplemented with 1 mg·mL^-1^ low endotoxin bovine serum albumin (BSA; ICPBio, Auckland, NZ; “assay medium”). Prior to any antibody labelling or drug treatment, cells were equilibrated in assay medium for 30 min at 37°C in 5% CO_2_.

Two immunocytochemical approaches for measuring the influence of ligands on receptor cell surface expression were utilised in this study (Fig. 1A). In “***Method A***”, cell surface receptors are labelled with primary antibody prior to drug treatment, and primary antibody remaining at the cell surface following drug treatment is quantified. Because only receptors that were at the cell surface prior to drug stimulation can be labelled, this method detects residual surface receptor expression, typically providing a direct measurement of receptor internalisation. Cells were incubated with mouse monoclonal anti-HA (901503; BioLegend, San Diego, CA, USA) diluted 1:500 in assay medium for 30 min at room temperature (RT). Cells were then washed with assay medium and stimulated with vehicle or drug for the indicated time period at 37°C. At the conclusion of the drug stimulation, plates were placed on ice for a minimum of 2 min to halt membrane trafficking and washed with RT assay medium. Alexa Fluor 488-conjugated goat anti-mouse secondary antibody (Invitrogen; diluted 1:300 in assay medium) was then applied to cells and incubated for 30 min at 15°C (preventing further trafficking [28,29]), prior to washing twice with assay medium and fixation (below).

**Fig. 1.**
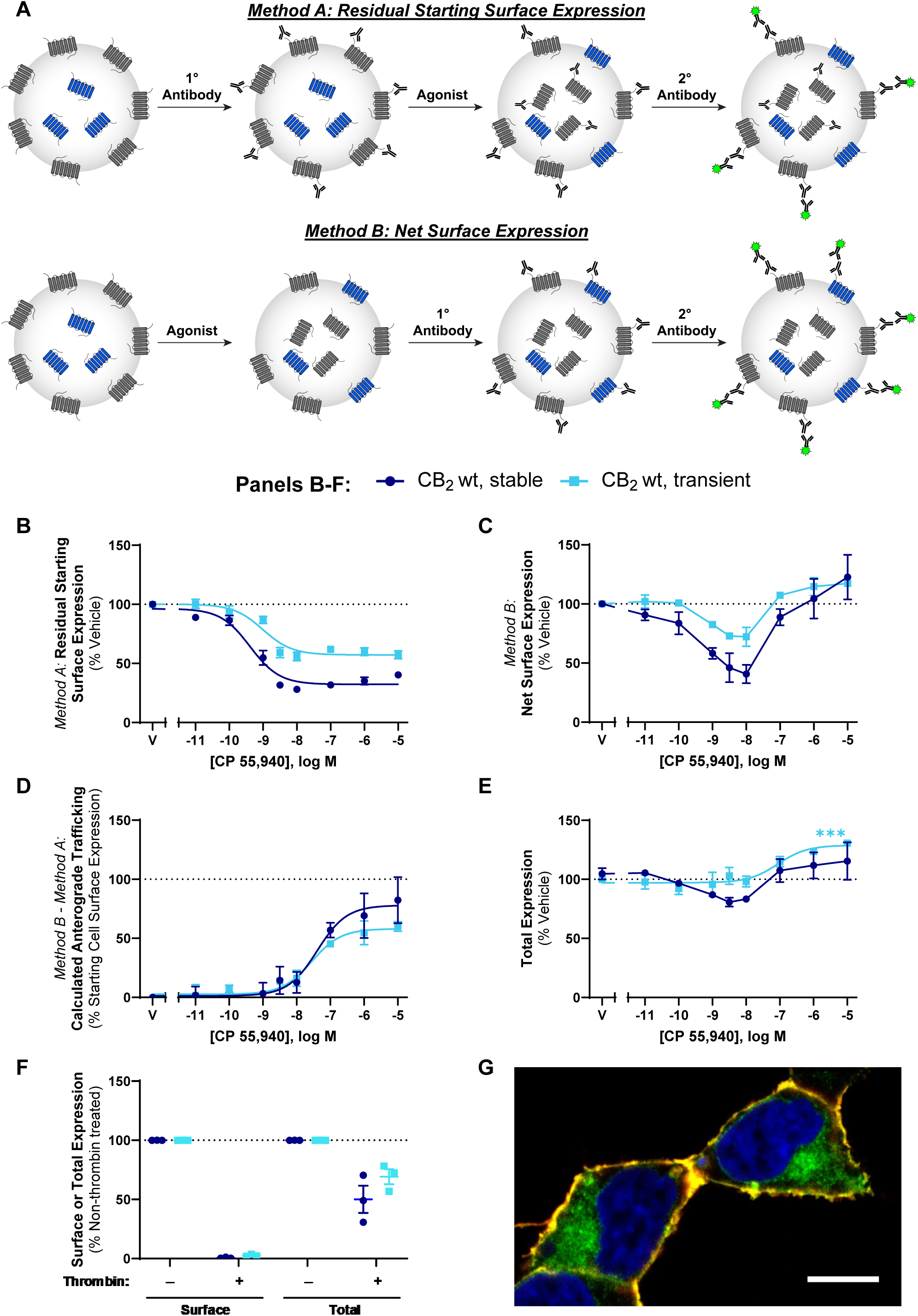
CB_2_ trafficking in response to CP 55,940. (A) Immunocytochemical approaches used to measure surface receptor expression. *Method A* – cells are labelled with primary antibody directed against an extracellular epitope tag on the receptor of interest, followed by stimulation with agonist. Cells are then incubated with secondary antibody under non-permeabilising conditions, to label residual primary antibody at the cell surface. *Method B* – cells are stimulated with agonist, then labelled with primary antibody to an extracellular epitope tag under non-permeabilising conditions. This labels receptors present at the cell surface at the end of drug stimulation. Cells are then fixed and secondary antibody applied. (B, C, E) HEK cells stably or transiently expressing hCB_2_ wt were treated with a CP 55,940 dilution series for 3 hours, and were labelled to measure (B) residual starting surface expression (*Method A*), (C) net surface expression (*Method B*), or (E) total receptor expression (i.e. primary antibody applied under permeabilising conditions to detect both cell surface and intracellular receptor). (D) Calculated anterograde trafficking (*Method B* – *Method A*). (F) Surface and total CB_2_ expression without and with exposure to thrombin (to cleave surface HA epitopes) prior to fixation. (B-F) Data are presented as mean ± SEM from three independent experiments. (G) Confocal image illustrating surface (red/yellow) and intracellular (green) CB_2_. Nuclei are stained blue. Scale bar, 10μm.

In contrast, “***Method B***” labels cell surface receptors with primary antibody *after* drug treatment. Therefore, as well as receptors that remain at the cell surface following drug stimulation, any receptors delivered to the cell surface during stimulation will also be detected. Cells were first stimulated with vehicle or drug for the indicated period of time at 37°C. At the conclusion of the drug stimulation, plates were placed on ice for a minimum of 2 min to halt membrane trafficking and washed with RT assay medium. Cells were then incubated with anti-HA diluted 1:500 in assay medium for 30 min at 15°C (preventing further trafficking), then washed twice with assay medium.

For both methods, cells were then fixed in 4% paraformaldehyde (PFA) at RT for 10 min and washed twice with phosphate buffered saline (PBS). Cells labelled via *Method B* were then incubated with Alexa Fluor 488-conjugated goat anti-mouse secondary (diluted 1:400 in immunobuffer: PBS with 0.2% Triton X-100 [PBS-T], 1% normal goat serum and 0.4 mg·mL^-1^ thiomersal [Merck, Darmstadt, Germany]), for 3 hours at RT or overnight at 4°C, and washed with PBS-T. For both methods, cell nuclei were then labelled with 8 µg·mL^-1^ Hoechst 33258 (Sigma-Aldrich) in PBS-T.

Where total receptor expression was measured, drug stimulations and cell fixation were carried out as described above but without any antibody labelling. When cell lines with stable receptor expression were assayed, 90% methanol was applied for 10 min at –20°C, then removed and the plate allowed to air dry for ∼3 min before re-hydrating in PBS-T. Cells were then incubated with anti-HA diluted 1:2500 in immunobuffer for 3 hours at RT or overnight at 4°C, washed with PBS-T, then labelled with secondary antibody and Hoechst as above.

For confocal imaging to illustrate surface versus intracellular CB_2_, cells were plated in chamber slides (BD Biosciences, Franklin Lakes, NJ, USA) and processed for immunocytochemistry, generally as above. In brief, live cells were cooled to 15°C and incubated with anti-HA then Alexa 488-conjugated secondary, fixed and permeabilised, then re-probed with anti-HA, followed by Alexa 594-conjugated secondary and Hoechst stain. Slides were mounted with AF1 antifadent (Citifluor, Hatfield, PA, USA).

#### 2.3.1 Cell surface receptor turnover

For assays measuring receptor turnover at the cell surface, cells were incubated with anti-HA primary antibody diluted 1:500 in assay medium for 30 min at RT. Cells were then stimulated with vehicle or 1 μM CP 55,940 for 3 hours at 37°C in the *continued* presence of anti-HA antibody. At the conclusion of the drug stimulation, cells were fixed in 4% PFA, incubated with Alexa Fluor 488-conjugated goat anti-mouse secondary in immunobuffer (with Triton X-100), and stained with Hoechst 33258 as above. The presence of anti-HA primary throughout the drug stimulation, and the application of secondary antibody under membrane permeabilising conditions, allowed the surface turnover of receptor to be detected (i.e. receptor that was delivered to and/or internalised from the surface at any point throughout the drug stimulation).

#### 2.3.2 Thrombin strip

In some experiments, proteolytic enzyme thrombin was used to cleave extracellular HA epitopes, thereby preventing labelling of surface HA-tagged receptor with anti-HA primary antibody but leaving intracellular HA epitopes intact. At the appropriate point in the assay, thrombin (50 U·mL^-1^; Bio-Pharm Inc., Hatfield, AR) or assay medium was applied to cells and incubated for 30 min at 37°C. Plates were placed on ice for a minimum of 2 min and washed with the protease inhibitor phenylmethylsulfonyl fluoride (PMSF, 250 µM; Sigma-Aldrich) to inactive thrombin. For experiments validating the effectiveness of the thrombin strip, cells were incubated with mouse monoclonal anti-HA diluted 1:500 in assay medium for 30 min at 15°C, and washed twice with assay medium to label any remaining HA epitopes at the cell surface. Cells were then fixed in 4% PFA. For experiments measuring intracellular versus total receptor (with and without thrombin strip, respectively), cells were methanol unmasked then anti-HA was incubated in immunobuffer as described above for total receptor detection. For both experimental designs, secondary antibody and Hoechst labelling was then carried out as described above.

#### 2.3.3 Inhibitors of signalling, trafficking, protein synthesis and monoacylglycerol lipase (MAGL)

For trafficking assays utilising PTX, cells were seeded in half the usual volume of medium. ∼5 hours later, PTX (final concentration 100 ng·mL^-1^) or vehicle were prepared in medium and applied to the pre-plated cells. Cells were exposed to PTX or vehicle (50 mM Tris pH 7.5, 10 mM glycine, 0.5M sodium chloride, 50% (v/v) glycerol; Sigma-Aldrich) for 16-20 hours prior to the start of the trafficking experiment.

Other inhibitors were pre-incubated at 2x their intended final concentration, after which CB_2_ agonist or vehicle was added which diluted the inhibitor to the 1x final concentration for the duration of the drug stimulation. Final concentrations and pre-incubation times were: Gallein 10 μM, 30 min; U73122 1 μM, 30 min; LY294002 10 μM, 30 min; GF109203X 1 μM, 30 min; BFA 1 µg·mL^-1^, 10 min; monensin 1 μM, 10 min; CHX 10 µg·mL^-1^, 10 min; JZL148 1 μM, 30 min.

### 2.4 Image acquisition and analysis

Assays utilising immunocytochemistry were quantified via automated image acquisition and analysis [6,30–32]. Briefly, images from four sites per well were captured using a 10x objective on an ImageXpress Micro XLS widefield microscope (Molecular Devices, Sunnyvale, CA) housed in the Biomedical Imaging Research Unit at the University of Auckland. Analysis was carried out with MetaXpress (v6.5, Molecular Devices) or MetaMorph (v7.10, Molecular Devices) software. User-defined thresholds were applied consistently within each experiment to define the boundary between genuine staining and background (“staining of interest threshold”), exclude brightly fluorescent debris if necessary, and to segment nuclei from background. The inbuilt ‘Find Spots’ algorithm was used to count the number of cells per image. Receptor expression was then quantified by integrated fluorescence intensity per cell [32]. For all trafficking assays, receptor expression was normalised to the vehicle-treated condition (100%). Anterograde trafficking was subsequently calculated by subtracting measurements for *Method A* from *Method B*.

Confocal images were taken on a Leica TCS SP2 system with 63x objective lens, as described previously [33].

### 2.5 Western Blotting

Cells were seeded in 24 well plates and transiently transfected with hCB_2_ wt, KK mutant, or benign plasmid [31] using Lipofectamine 2000 (Thermo Fisher Scientific). Drugs and reagents were prepared, and cells stimulated 18-24 hours post-transfection, as described in Section 2.3 for 3 hours. Plates were then placed on ice for 2 min and washed with RT assay medium.

Cells were harvested in lysis buffer (150 mM NaCl, 1% Triton X-100, 50 mM Tris-HCl pH 8, cOmplete protease inhibitor cocktail tablets [Roche]), incubated on ice for 30 min and centrifuged for 10 min at 4°C. Supernatants were aspirated and diluted with 4x loading buffer (125 mM Tris-HCl pH 6.8, 50% glycerol, 4% sodium dodecyl sulfate [SDS]), and protein concentrations quantified. Cell lysate concentrations were standardised in 1x loading buffer with 0.001% bromophenol blue and heated for 30 min at 37°C. Lysates (15 μg per well) were separated on a 4-12% pre-cast gel (Invitrogen) by electrophoresis, followed by semi-dry transfer to polyvinylidene fluoride membranes. Membranes were air dried and rehydrated in methanol, rinsed, and blocked in 5% non-fat milk (NFM) in 0.1 M Tris-buffered saline with 0.2% Triton X-100 (TBS-T) for 30 min at RT. Mouse monoclonal anti-HA (901503; BioLegend) and rabbit polyclonal anti-glyceraldehyde 3-phosphate dehydrogenase (GAPDH; ab9485; Abcam) diluted 1:1000 in 1% NFM/TBS-T were co-incubated for 3 hours at RT or overnight at 4°C. Following three TBS-T washes, membranes were co-incubated with fluorescently conjugated secondary antibodies (IRDye^®^ 680LT goat anti-mouse [926-68020; LI-COR Biosciences, Lincoln, NE] and IRDye^®^ 800CW goat anti-rabbit [926-32211; LI-COR Biosciences]) diluted 1:20 000 in 1% NFM/0.01% SDS/TBS-T for 3 hours at RT or overnight at 4°C. Membranes were then rinsed and left to air dry prior to imaging.

Images were captured on an Odyssey^®^ FC Imaging System (LI-COR Biosciences) in 700 nm and 800 nm channels, and quantified using Image Studio (Lite; LI-Cor Biosciences). GAPDH and HA bands were segmented using the shapes tool and integrated intensity above background obtained. In the anti-HA channel, the corresponding HEK wt (i.e. no CB_2_) measurement was subtracted from the total signal for each band/range of bands, and the remaining signal normalised to its respective GAPDH. Within each treatment condition (i.e. lane), signals for each distinct band or range of molecular weights, were then normalised to the total signal in the lane for that particular treatment condition to determine what proportion each band, or range of molecular weights, contributed to total expression. Uncropped blot images were provided for peer review.

### 2.6 General analysis and statistics

All graphs were prepared in GraphPad Prism (v10; GraphPad Software Inc., La Jolla, CA). Sigmoidal three-parameter nonlinear regression concentration-response curves were fitted and parameters obtained (“span” and logEC_50_) for independent experiments. Statistical analyses were performed on data from three to four independent experiments using Sigmaplot™ v13.0 (Systat Software Inc., San Jose, CA, USA). The Shapiro-Wilk test for normality and Brown-Forsythe test for equal variance were performed to verify the datasets were appropriate for analysis with parametric statistical tests. Where a set of results did not pass either of these tests, datasets were transformed (log_10_ or *e^x^*) to change the nominal values of the sample distribution, and re-tested. Following a pass result of *p* > 0.05, a Student’s t-test, one-way ANOVA, or two-way ANOVA was carried out as appropriate for the number of conditions and factors under comparison. A paired or repeated measures design was used where appropriate. If a statistically significant difference (*p* < 0.05) was detected, the Holm-Šídák [34] post-hoc test was used to test for significant differences within/between groups. *P* values are indicated graphically as: * < 0.05, ** < 0.005, *** < 0.001.

### 2.7 Radioligand binding

Radioligand binding was carried out as in [35], with minor modifications and following a saturation binding assay design. Cell membranes were prepared from HEK cells stably expressing CB_2_ wt (also used for trafficking experiments) as described previously [36], and concentration determined by protein assay (“DC”, Bio-Rad, CA, USA). Membranes (10 μg per point) were incubated in a total volume of 200 uL at 30°C for 1 hour with 1-20 nM ^3^H-CP 55,940 (Perkin Elmer, MA, USA) and *either* 1 μM CP 55,940 (non-specific binding) or vehicle (total binding) in binding buffer (50 mM HEPES pH 7.4, 1 mM MgCl_2_, 1 mM CaCl^2^, 2 mg·mL^-1^ BSA). After incubation, samples were applied under vacuum to a 96-well GF/C glass fibre plate (Perkin Elmer) that had been pre-treated with 0.1% polyethylenimine (Sigma-Aldrich), followed by three rapid washes with ice cold wash buffer (50 mM HEPES pH 7.4, 500 mM NaCl, 1 mg·mL^-1^ BSA). After drying overnight, 50 μL per well Irgasafe Plus (Perkin Elmer) scintillation fluid was applied and plates were read in a MicroBeta TriLux 1450 scintillation counter (Perkin Elmer) for 1 min per well. Data were analysed in GraphPad Prism by “one site (total and nonspecific binding)” nonlinear regression, with 1/Y^2^ weighting and “Background” constrained to 0. For each assay, maximum specific binding (Bmax) in corrected counts per minute (ccpm) was converted to fmol·mg^-1^ via the manufacturer-supplied radioligand specific activity (141.2 Ci·mmol^-1^, 313.5 dpm·fmol^-1^) after adjusting for expected decay at time of assay based on tritium half-life (∼96% activity remaining). CB_2_ expression in fmol·mg^-1^ is provided as mean ± SEM from three independent experiments.

### 2.8 cAMP (CAMYEL biosensor)

Cyclic AMP (cAMP) assays were carried out utilising the CAMYEL biosensor (His-CAMYEL in pcDNA3L; ATCC MBA-277), as described previously [31,37] with minor modifications. HEK cells stably expressing CB_2_ wt (also used for trafficking experiments), or the untransfected parental line, were seeded at 36,000 cells/well in PDL-treated white 96-well plates (Corning) supplemented with 15% FBS. The same day, 24 ng/well CAMYEL plasmid was incubated with 0.25 μg/well PEI Max (Polysciences, PA, USA) in DMEM (without serum) for 60 min at room temperature, then added to plated cells (thereby adjusting FBS concentration to 10%). After incubating for ∼48 hours, cells were washed with and equilibrated in assay buffer (Hank’s Balanced Salt Solution [Thermo Fisher Scientific] with 1 mg·mL^-1^ BSA) for 30 min at 37°C. Coelenterazine h (Nanolight Technology, AZ, USA) 5 μM was applied to cells, and baseline BRET measurements taken at ∼1 min intervals for ∼5 min in a CLARIOstar plate reader (BMG Labtech, Ortenberg, Germany; Rluc and YFP detected via monochromator with emissions bands 470 ± 30 nm and 555 ± 45 nm, respectively) at 37°C. A range of concentrations of Δ9-THC or CP 55,940 (keeping vehicle concentration constant for all) with forskolin (final concentration on cells 5 μM; Tocris Bioscience) were then added, and BRET measurements taken. BRET ratios (555 / 470 nm) were calculated, and the mean ratios for 0-10 min of drug incubation taken. Data were then normalised to the forskolin (with vehicle; 100%) and vehicle-only (0%) controls, and results from three independent experiments represented as means ± SEM.

## 3 Results

### 3.1 Agonist stimulation induces concurrent CB_2_ internalisation and anterograde trafficking

We aimed to extend our understanding of CB_2_ intracellular trafficking, subcellular distribution and post-endocytic fate utilising different immunocytochemistry paradigms to detect cell surface CB_2_. In pilot experiments, we observed a method-dependent difference in results which was not evident within the time period usually associated with internalisation (∼30 minutes) but was clear after a 3 hour stimulation and was hence explored further. The effect of a 3 hour stimulation with CP 55,940 on cell surface expression of both stably– and transiently-expressed hCB_2_ wt was investigated via two immunocytochemical labelling methods that we refer to herein as *Method A* and *Method B* (Fig. 1A). When surface CB_2_ was labelled with antibody *prior to* agonist stimulation (*Method A*), an agonist concentration-dependent decrease in CB_2_ surface expression was observed, which is indicative of agonist-induced internalisation (Fig. 1B, Table 1). Potency was approximately equivalent between stable and transient CB_2_ expression (*p* > 0.05). However, the proportional extent of internalisation was greater in the stably-expressing cells, whereby ∼20% more of the starting surface population was internalised (*p =* 0.004). Our stable cell line expressed CB_2_ at 2100 fmol·mg^-1^ (± 66, radioligand binding assay), approximately twice that of mixed human primary peripheral blood mononuclear cells in a prior report [38]. Transiently-expressing cells had 8.6 (±1.3) times the surface expression and 101 (±29) times the total expression of our stable cell line (by immunocytochemistry). The lesser internalisation extent in transiently-expressing cells was therefore likely a consequence of the greater surface (and overall) CB_2_ expression level in the transiently-expressing cells, and consistent with a prior report [9].

**Table 1.** Summary data for stably– and transiently-expressed CB_2_ trafficking in response to CP 55,940. Parameters are derived from data shown in Fig. 1B and 1D, from three independent experiments.

|  | *Method A: Internalisation (3 hour)* |  | *Method B – Method A:*<br>*Calculated Anterograde Trafficking*<br>*(3 hour)* |  |
| --- | --- | --- | --- | --- |
|  | pEC<sub>50</sub> (± SEM) | E<sub>max</sub>, % Vehicle<br>(± SEM) | pEC<sub>50</sub> (± SEM) | E<sub>max</sub>, % Starting<br>Surface Expression,<br>Vehicle (± SEM) |
| **Stable CB<sub>2</sub> wt** | 9.40 (0.09) | -64.4 (2.4) | 7.35 (0.17) | 76.6 (10.0) |
| **Transient CB<sub>2</sub> wt** | 9.09 (0.19) | -43.4 (2.4) | 7.66 (0.29) | 56.7 (5.5) |

When surface CB_2_ was labelled *following* CP 55,940 stimulation via *Method B* (monitoring net surface expression), surface CB_2_ expression was reduced by CP 55,940 concentrations in the region of the EC_50_ for internalisation (Fig. 1C). Conversely, at greater concentrations a lesser, or lack of, change in surface CB_2_ was observed in both expression conditions. Unlike *Method A*, which measures the proportion of initial surface CB_2_ remaining, *Method B* is a measure of net surface expression. It is apparent from Fig. 1B that the receptors initially on the cell surface and internalised by CP 55,940 remain internalised. A prior study utilising *Method A* has validated that the antibody used to label cell surface CB_2_ remains bound to the receptor, with no detectable degradation after a 4 hour agonist stimulation, and can be detected to recycle when agonist is replaced by inverse agonist [6]. Therefore, plain interpretation of the discrepancy between the findings from labelling *Methods A* and *B* (Fig. 1B and C) is that, despite cell surface CB_2_ undergoing internalisation, CP 55,940 can *also* stimulate delivery of non-cell surface-derived CB_2_ to the plasma membrane. We hypothesised that the source of the delivered receptors may be the “intracellular pool” described in earlier studies. Indeed, in our cells approximately 50-70% of overall CB_2_ expression was observed to be intracellular, as determined by comparing total CB_2_ expression with and without enzymatic cleavage of the surface HA tag, and intracellular expression was also evident in confocal imaging (Fig. 1F and G).

A reduction in surface expression in *Method A* is indicative of receptor internalisation, whereas *Method B* measures net surface expression which is a product of both internalisation and delivery of new receptors to the cell surface. The difference between these methods therefore reflects stimulated anterograde delivery of CB_2_ to the cell surface (Fig. 1D, Table 1). The potency of CP 55,940-induced CB_2_ anterograde trafficking at a 3 hour time point, was approximately 30-to 100-fold lower than that for internalisation. The maximal extent of stimulated cell surface re-population was equivalent to around 60-80% of cell surface expression in unstimulated cells (Fig. 1D, Table 1).

Total cellular CB_2_ expression was also monitored following CP 55,940 stimulation. This revealed increases in expression at CP 55,940 concentrations that induced anterograde trafficking of CB_2_ in both expression conditions (Fig. 1E). The maximal increase in expression measured was only significantly different from vehicle when hCB_2_ wt was transiently expressed (Fig. 1E; *p <* 0.001). In addition, slight decreases in total expression were observed at intermediate CP 55,940 concentrations in the stably-expressing line, but not with transient hCB_2_ wt expression.

Despite transient expression producing greater overall CB_2_ expression per cell than stable expression, findings between stably– and transiently-expressed CB_2_ in Fig. 1 were largely similar, validating the use of both transiently– and stably-expressing cells for further characterisation of these processes.

To further investigate the time-dependence of the CB_2_ trafficking phenotype, a time course over 6 hours was carried out in response to 1 μM CP 55,940 (a concentration that induced a pronounced degree of anterograde trafficking at 3 hours). When *Method A* was used, the application of primary antibody was staggered so that constitutive internalisation could be measured over the time course, as seen in the “vehicle” condition of Fig. 2A. Approximately 45% of CB_2_ was constitutively internalised over the 6 hours, consistent with our prior findings [6]. Approximately 70% of surface CB_2_ internalised in response to CP 55,940 within 0.5 hours, and the residual surface expression remained unchanged over the remaining time course. When *Method B* was used, a reduction relative to vehicle was observed at 0.5 and 1 hour, followed by no change and/or an increase at later time points (Fig. 2B). Subtraction of *Method A* from *Method B* to reveal any anterograde trafficking/surface delivery of hCB_2_ wt allowed measurement of constitutive surface delivery in the vehicle condition, and the combination of constitutive and ligand-induced surface delivery in the CP 55,940 condition (Fig. 2C). The difference between the vehicle and CP 55,940 traces therefore corresponds to CP 55,940-induced surface delivery. Anterograde trafficking of hCB_2_ wt in response to CP 55,940 was significantly different from vehicle at all time points > 1 hour (*p =* 0.005, < 0.001, < 0.001 respectively), indicating that this phenotype is detectable following a lag time of at least 1 hour. CP 55,940-induced surface delivery continued at a steady rate over the remainder of the time course.

**Fig. 2.**
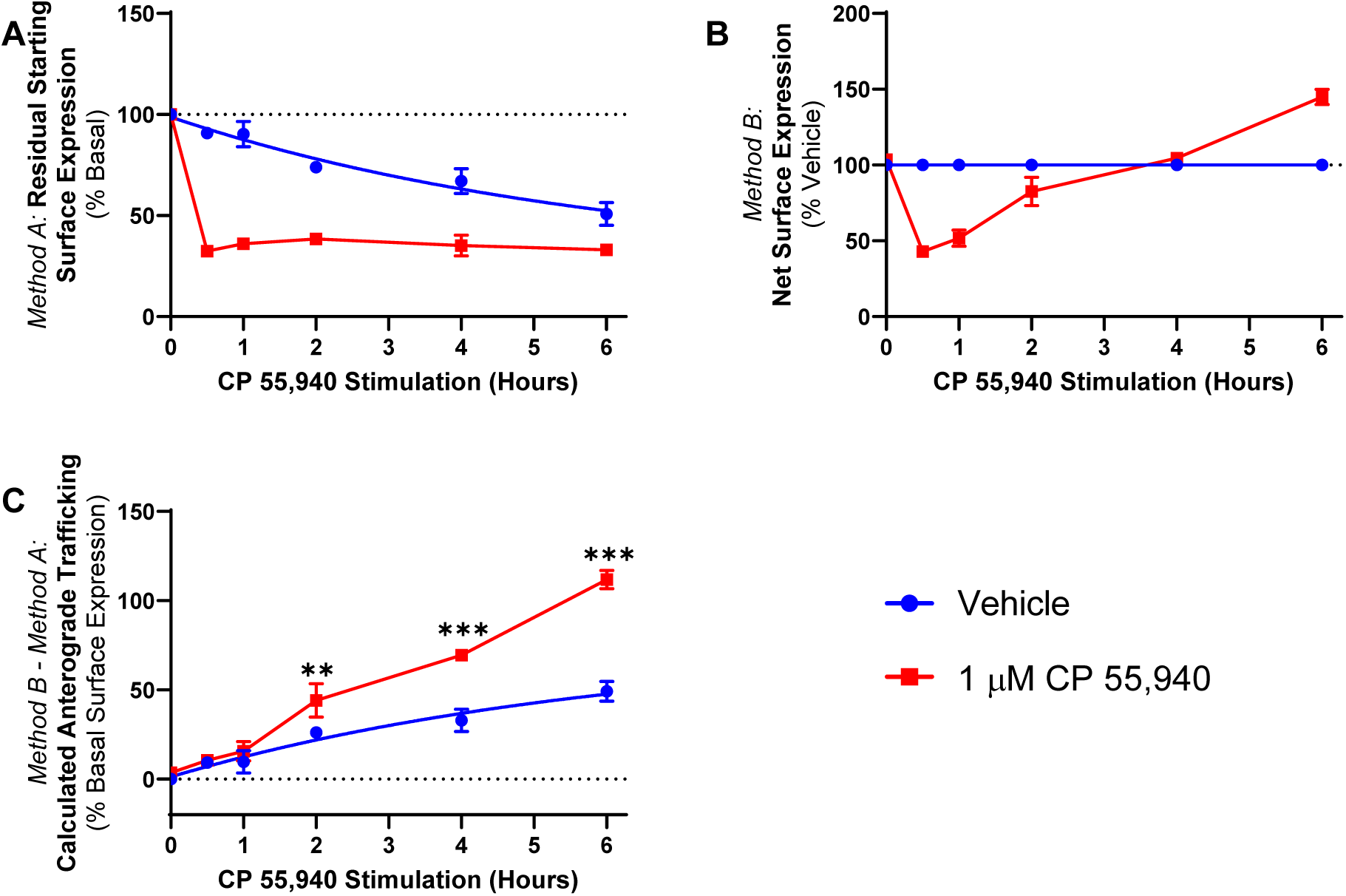
CB_2_ trafficking time course in response to CP 55,940 HEK cells stably expressing CB_2_ wt were treated with 1 µM CP 55,940 or vehicle over a 6 h time course, and labelled to measure. (A) residual starting surface expression (*Method A*) or (B) net surface expression (*Method B*). (C) Calculated anterograde trafficking (*Method B* – *Method A*). *P* values for significant differences between Vehicle and CP 55,940 are indicated graphically as: ** < 0.005, *** < 0.001. Data are presented as mean ± SEM from three independent experiments.

To explore whether CP 55,940-induced anterograde trafficking was particular to CB_2_, concentration responses using the labelling methods described above were carried out in HEK cells stably expressing HA-tagged CB_1_ or hβ_2_-AR. As CP 55,940 is a full agonist at CB_1_ (in addition to CB_2_), and β_2_-AR is a well-characterised recycling receptor, these receptors are appropriate controls in this context. The β_2_-AR-specific ligands salbutamol and isoproterenol induced a concentration-dependent decrease in surface hβ_2_-AR expression, and CB_1_ was similarly internalised in response to CP 55,940 (Fig. 3A and B). In both cases, the two labelling methods produced interchangeable concentration response curves with no evidence of ligand-stimulated anterograde trafficking at any ligand concentration. In addition, when CP 55,940 was applied to β_2_-AR at concentrations shown to either internalise or stimulate surface delivery of hCB_2_ wt, (i.e. 10 nM and 1 µM, respectively), surface receptor expression was equivalent between labelling conditions (Fig. 3C). This indicates that CP 55,940-induced anterograde trafficking is CB_2_-specific, and not an artefact of the labelling methods employed or a result of a general effect on receptor trafficking in the HEK cell line, nor a result of modified control of the constitutive CMV promoter (as hβ_2_-AR and hCB_2_ were both expressed under the CMV promoter).

**Fig. 3.**
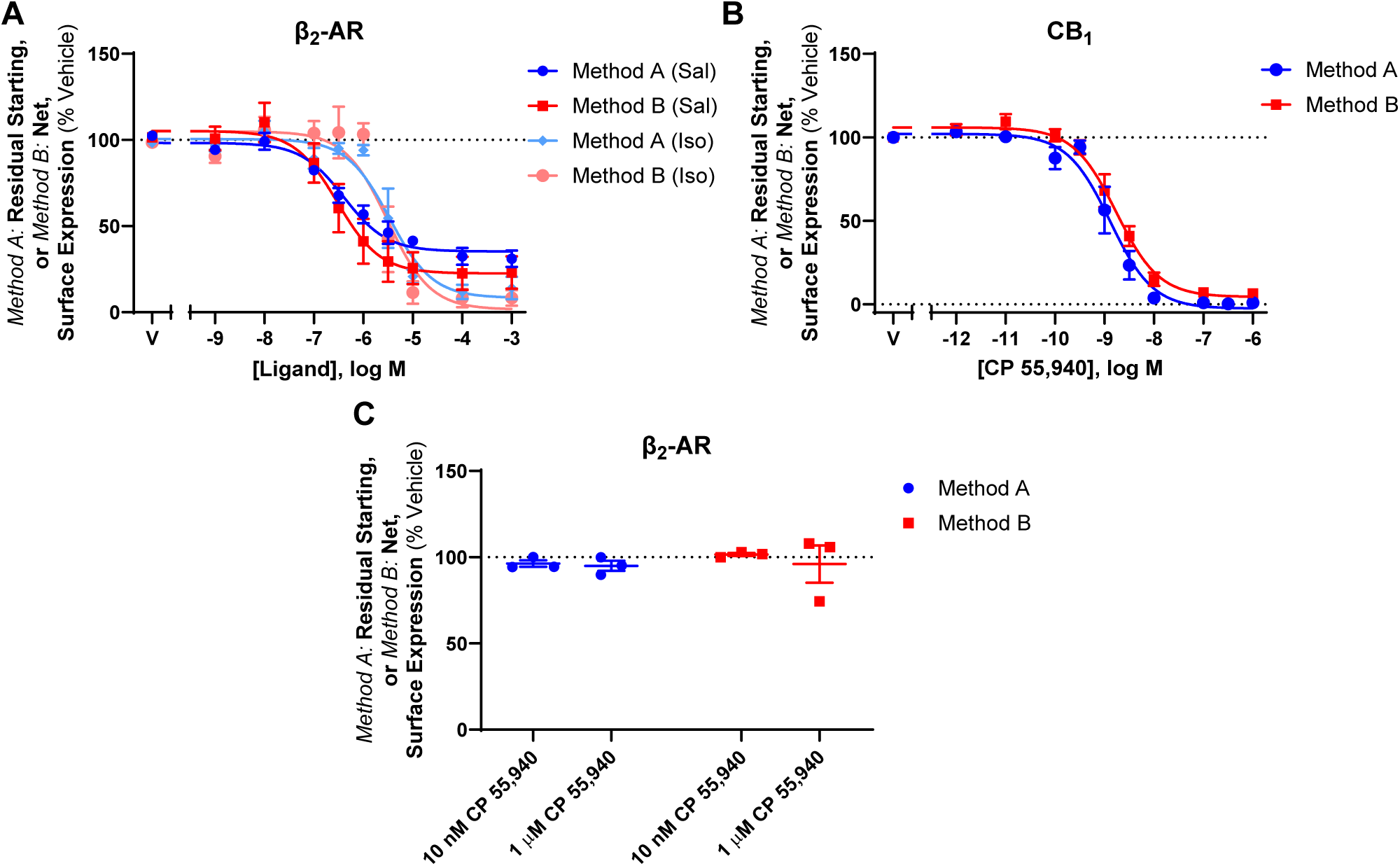
Trafficking of β_2_-AR and CB_1_ HEK cells stably expressing β_2_-AR were treated with. (A) a Salbutamol (Sal) or Isoproterenol (Iso) dilution series, or (C) an internalising (10 nM) or anterograde trafficking-inducing (1 µM) concentration of CP 55,940 for 3 h. (B) HEK cells stably expressing CB_1_ were treated with a CP 55,940 dilution series for 3 h. Residual starting surface expression (*Method A*) and net surface expression (*Method B*) were measured in all assays. Data are presented as mean ± SEM from three independent experiments.

### 3.2 A di-lysine ‘KK’ motif is required for basal surface delivery of CB_2_ but not CP 55,940-induced anterograde trafficking

CB_2_ has *ExE* and *KK* motifs in its cytoplasmic (C-terminal) tail, which have been shown to modulate secretory pathway trafficking in other GPCRs [39–42] and non-GPCR membrane proteins [43,44][45]. CB_2_ constructs (Fig. 4A) containing mutations to these motifs (ETE339-41ATA and KK318-9EE), as well as a truncated Δ12 mutant (lacking the last 12 residues of the CB_2_ C-terminal tail, 349-360, including residues that may influence trafficking: S352 phosphorylation site [5], a cluster of aspartic acid residues that may act a phospho-mimetics [46], and a type 2 PDZ-binding motif [47], were screened to determine their effects on internalisation and net surface expression in response to CP 55,940 relative hCB_2_ wt. The ETE and Δ12 mutants had similar basal surface expression and trafficked equivalently to wt] in response to CP 55,940 (Fig. 4B and C). In contrast, mutation of the KK motif abolished basal surface expression of CB_2_, indicating that this motif influences typical receptor trafficking or synthesis. Despite this, exposure to CP 55,940 stimulated surface expression of the KK mutant. Total expression levels of the ETE and KK mutants were not significantly different from hCB_2_ wt, indicating that none of these mutations negatively affected overall receptor synthesis and/or turnover (Fig. 4D). However, deletion of the last 12 residues of the CB_2_ C-terminal tail significantly increased total expression (*p* = 0.005 for Δ12 vs. CB_2_ wt).

**Fig. 4.**
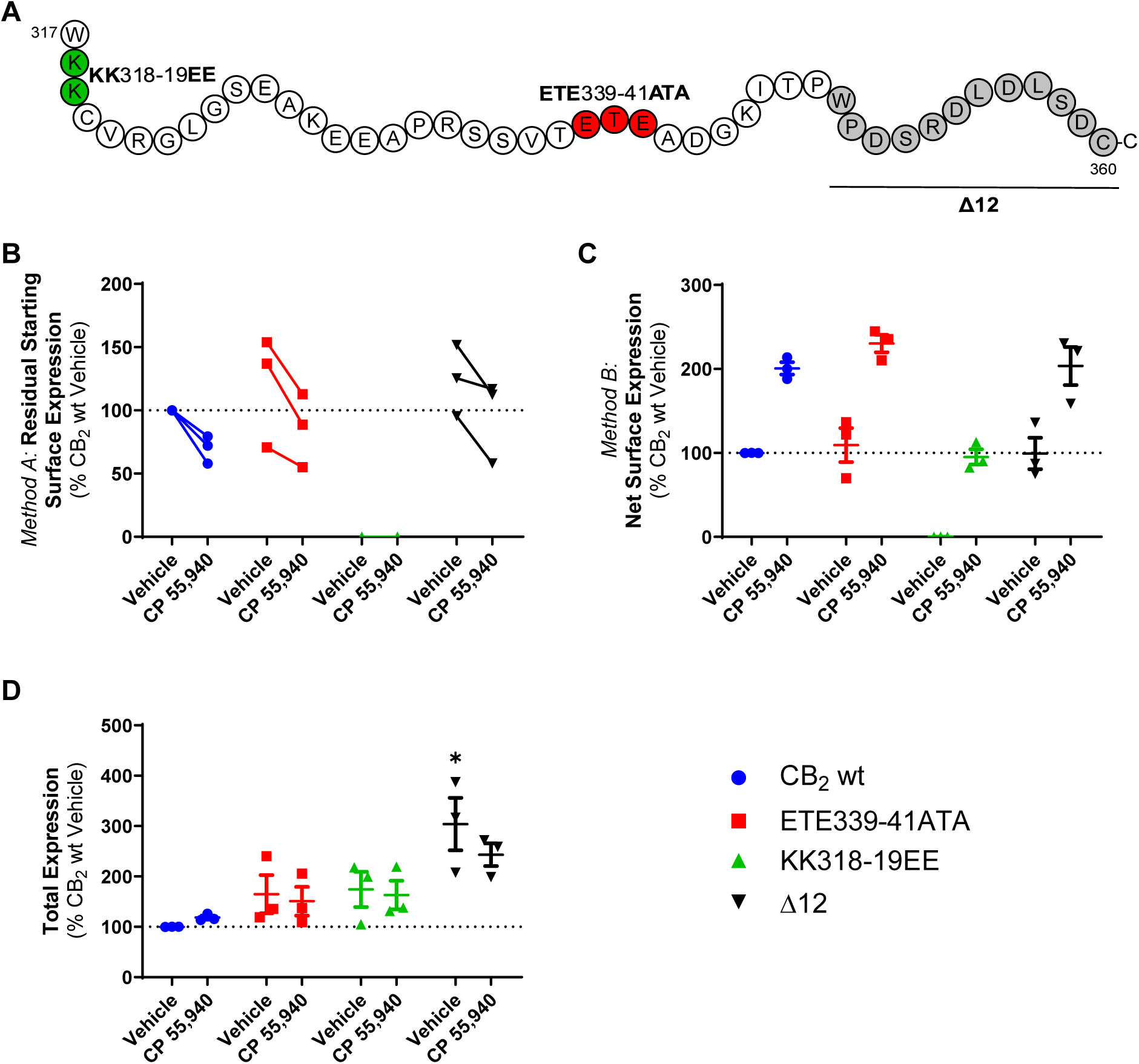
Trafficking of CB_2_ mutant constructs in response to CP 55,940 HEK cells transiently expressing CB_2_ wt or CB_2_ C-terminal tail mutants. (A) were treated with vehicle or 1 μM CP 55,940 for 3 h and labelled to measure (B) residual starting surface expression (*Method A*), (C) net surface expression (*Method B*), or (D) total receptor expression. Data are presented as mean ± SEM from three independent experiments, except in Panel B where matched data from each of three independent experiments is plotted. *P* values for significant differences between CB_2_ wt and mutant vehicle conditions are indicated graphically as: * < 0.05.

Assays measuring the effect of CP 55,940 on the surface expression of transiently and stably expressed KK mutant receptor were performed (analogous to those for hCB_2_ wt in Fig. 1). As expected, no basal surface expression of the KK mutant was detected when receptor was labelled prior to agonist stimulation (*Method A*, Fig. 5A). However, a CP 55,940 concentration-dependent increase in net surface expression was observed, and this was equivalent between transfection conditions (Fig. 5B). The ultimate extents of CP 55,940 induced-surface delivery were equivalent between hCB_2_ wt and KK mutant, but the effect was ∼1 log unit more potent for hCB_2_ wt (Fig. 1D versus 5C; Table 1 versus Table 2). As with hCB_2_ wt, total expression of the KK mutant increased at CP 55,940 concentrations that also induced anterograde trafficking (Fig. 5D).

**Fig. 5.**
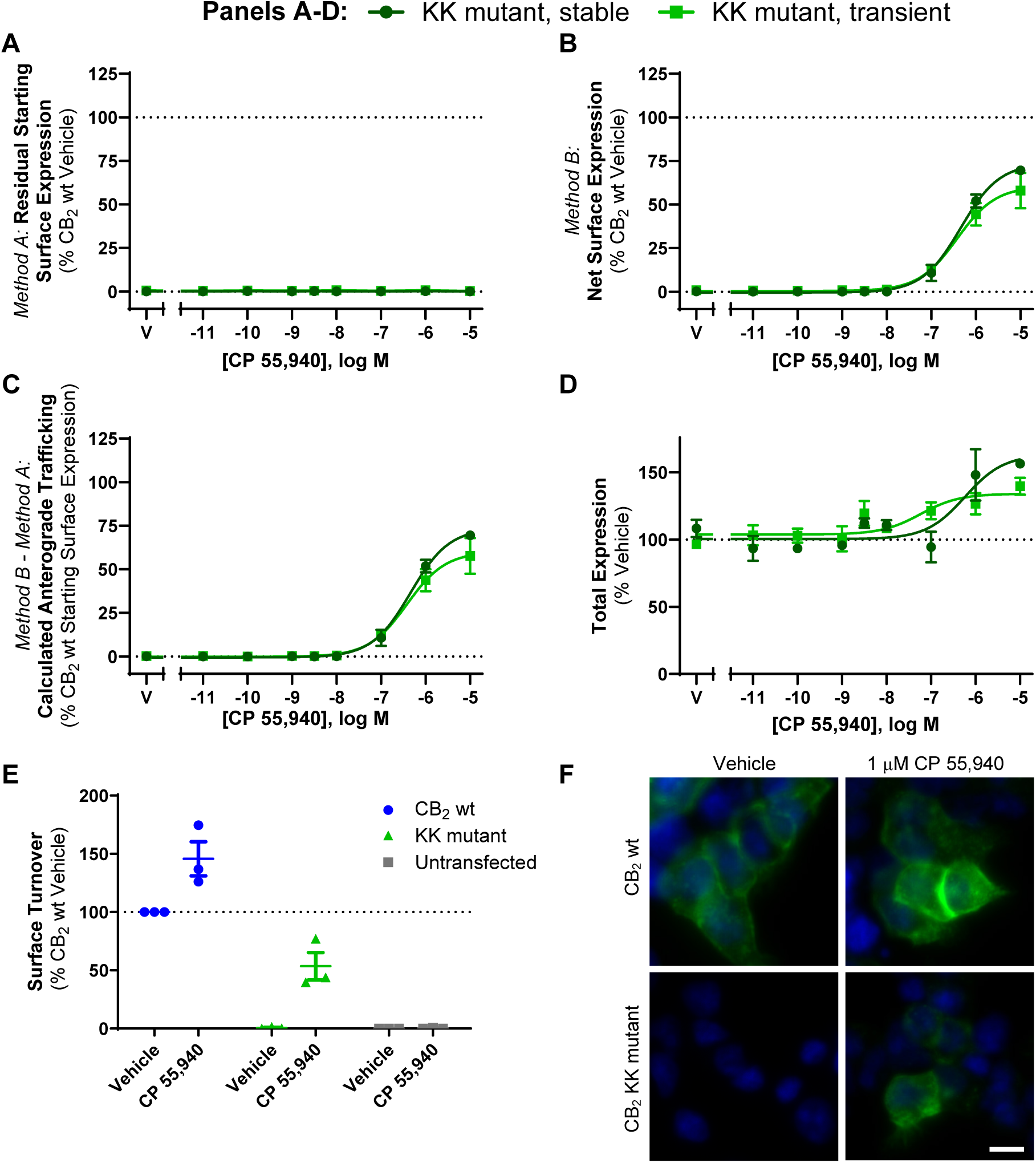
CB_2_ KK mutant trafficking and expression in response to CP 55,940 HEK cells stably and transiently expressing KK mutant were treated with a CP 55,940 dilution series for 3 h and were labelled to measure. (A) residual starting surface expression (*Method A*), (B) net surface expression (*Method B*), or (D) total receptor expression. (C) Calculated anterograde trafficking (*Method B* – *Method A*). Data are presented as mean ± SEM from three independent experiments. (E-F) HEK cells transiently expressing CB_2_ wt or KK mutant were treated with vehicle or 1 μM CP 55,940 for 3 h in the continued presence of primary anti-HA to measure surface turnover. (E) Mean ± SEM from three independent experiments. (F) CB_2_ (anti-HA) in green, nuclei (Hoechst stain) in blue. Scale bar = 10 μM. Representative of three independent experiments.

**Table 2.** Summary data for stably– and transiently-expressed CB_2_ KK mutant trafficking in response to CP 55,940. Parameters are derived from data shown in Fig. 5C, from three independent experiments.

|  | *Method B – Method A:*<br>*Calculated Anterograde Trafficking*<br>*(3 hour)* |  |
| --- | --- | --- |
|  | pEC<sub>50</sub> (± SEM) | E<sub>max</sub>, % CB<sub>2</sub> wt<br>Vehicle (± SEM) |
| **Stable CB<sub>2</sub> KK mutant** | 6.35 (0.11) | 74.3 (2.7) |
| **Transient CB<sub>2</sub> KK mutant** | 6.42 (0.11) | 60.86 (10.2) |

Lack of basal surface expression of the KK mutant could be due to either impaired delivery to the cell surface, or efficient surface delivery followed by rapid constitutive internalisation. To clarify this, we measured surface turnover of hCB_2_ wt versus the KK mutant during a 3 hour incubation in media (with vehicle) or CP 55,940 (Fig. 5E-F). “Surface turnover” was assessed by incubating cells in the continued presence of anti-HA, such that receptor trafficking via the cell surface at any point during the assay would bind antibody. Secondary antibody was then applied under permeabilising conditions to detect anti-HA bound receptor anywhere in the cell, even if it had subsequently internalised. Stimulation of hCB_2_ wt with CP 55,940 enhanced surface turnover compared with vehicle, consistent with CP 55,940-induced surface delivery. The equivalent was true for the KK mutant, which had a considerable amount of antibody bound in response to CP 55,940 stimulation. In contrast, no antibody labelling could be detected for KK mutant-expressing cells incubated with vehicle only, indicating lack of constitutive receptor delivery to the cell surface over 3 hours (Fig. 5E and F). Non-specific antibody uptake (in untransfected cells) was below detectable limits (Fig. 5E). Combined with previous findings, this indicates that in unstimulated cells the KK318-9EE mutation prevents constitutive CB_2_ delivery to the plasma membrane.

### 3.3 hCB2 wt and KK mutant possess the same maturation states in differing proportions, with maturation states altered by agonist stimulation

From this data, as well as prior findings in other receptors [39,40,42], we hypothesised that the KK318-9EE mutation might inhibit hCB_2_ from traversing and/or exiting the secretory pathway basally, whereas stimulation with CP 55,940 seems to release this blockade. Western blotting revealed numerous molecular weight species in both CB_2_ wt and KK mutant lysates, of which the three most prominent and size-resolved bands were ∼35 kDa, ∼40 kDa, and ∼67 kDa, with species of varying size also evident between ∼40 and ∼55 kDa, and above ∼67 kDa (Fig. 6A). As the predicted molecular weight of unmodified CB_2_ wt and KK mutant constructs is ∼42.5 kDa, the ∼35 kDa species likely represents immature CB_2_. Indeed, receptor species have previously been observed to be smaller than their predicted size via western blot [33,48,49]. The ∼40 kDa species observed here is therefore likely to be a more mature form of CB_2_, based on a prior report of a 5 kDa difference between mature and de-glycosylated human CB_2_ [50], while the >40 to ∼55 kDa forms might have more complex glycosylation or other post-translational modifications, and/or be complexed with small interacting proteins. The ∼67 kDa and larger species might reflect CB_2_ homodimer (previously shown to be ∼87 kDa in [50]) or CB_2_ complexed with other interacting protein(s). As expected, no anti-HA staining was detected in the HEK wt control, indicating specificity for HA-tagged CB_2_.

**Fig. 6.**
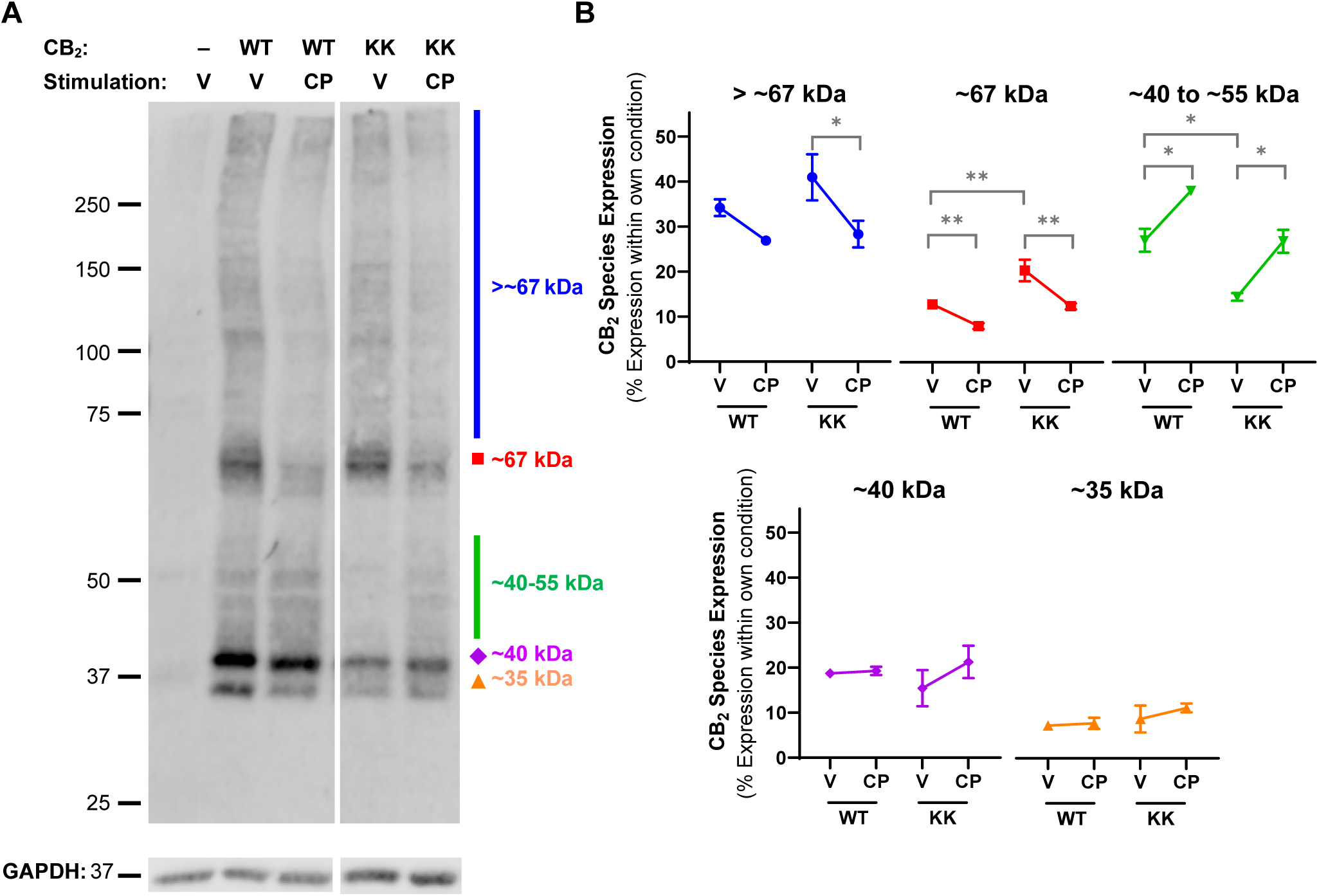
Western blot analysis for CB_2_ wt and KK mutant HEK cells transiently expressing CB_2_ wt (“WT”), KK mutant (“KK”), or benign vector, were treated with vehicle (V) or 1 µM CP 55,940 (CP) for 3 hours. (A) Western blot for anti-HA to detect CB_2_ (top panel) or GAPDH control protein (bottom panel). Molecular weight markers shown in kDa. All lanes are from the same membrane and displayed at the same intensity, but re-arranged for presentation. Representative of three independent experiments. (B) Each distinct band, or molecular weight range, was normalised to the total signal from the entire lane within the same experimental condition – i.e. each is represented as a percentage of total receptor expression. *P* values for significant differences between Vehicle and CP 55,940 or WT and KK (comparisons indicated by brackets) are indicated graphically as: * <0.05, ** < 0.005. Data are presented as mean ± SEM of three independent experiments.

Blots were analysed quantitatively for proportional expression of molecular weight species within each receptor type and treatment condition (Fig. 6B). The smallest ∼35 and ∼40 kDa bands represented fairly equivalent proportions of total receptor expression between wt and KK mutant CB_2_, and were unchanged by CP 55,940. The >67 kDa and ∼67 kDa species made up a larger proportion of KK mutant relative to CB_2_ wt in vehicle-treated cells (∼41% versus ∼34% and ∼20% versus 12%, respectively), though only the latter was significantly different (p = 0.001). Conversely, species ranging from ∼40 to 55 kDa were more prominent in CB_2_ wt than in the KK mutant, comprising ∼27% of total CB_2_ wt expression, but ∼15% of KK mutant (p = 0.021). CP 55,940 stimulation appeared to reduce the proportional expression of the ≥67 kDa species for both WT and KK mutant, with significant differences for all (p = 0.001 – 0.032) except CB_2_ wt >67 kDa species. In an inverse shift, the proportional expression of species ranging from ∼40 to 55 kDa increased in response to CP 55,940 for both receptors (p = 0.017 for CB_2_ wt, p = 0.027 for KK mutant).

### 3.4 CP 55,940-induced anterograde trafficking of CB_2_ does not require common signalling pathways

We hypothesised that a mechanism involving CB_2_ signalling may be a feedback mechanism stimulating the CP 55,940-induced CB_2_ cell surface delivery observed. To test this, we measured residual starting surface and net surface expression concentration response curves in response to CP 55,940 in the absence and presence of various signalling inhibitors: irreversible Gα_i/o_ and Gβγ inhibitor PTX, Gβγ inhibitor gallein, phospholipase C [PLC] inhibitor U73122, phosphoinositide 3-kinase [PI3K] inhibitor LY249002, and protein kinase C [PKC] inhibitor GF109203X.

CP 55,940-stimulated CB_2_ internalisation potency and efficacy were unaffected by the signalling inhibitors (*p* > 0.05; Fig. 7A and Table 3). The signalling inhibitors had no significant effects on CP 55,940-stimulated surface delivery of hCB_2_ wt or KK mutant (*p* > 0.05; Fig. 7C-D, Table 3). Cell morphology and number were also unaffected by signalling inhibitors (Fig. 7G).

**Fig. 7.**
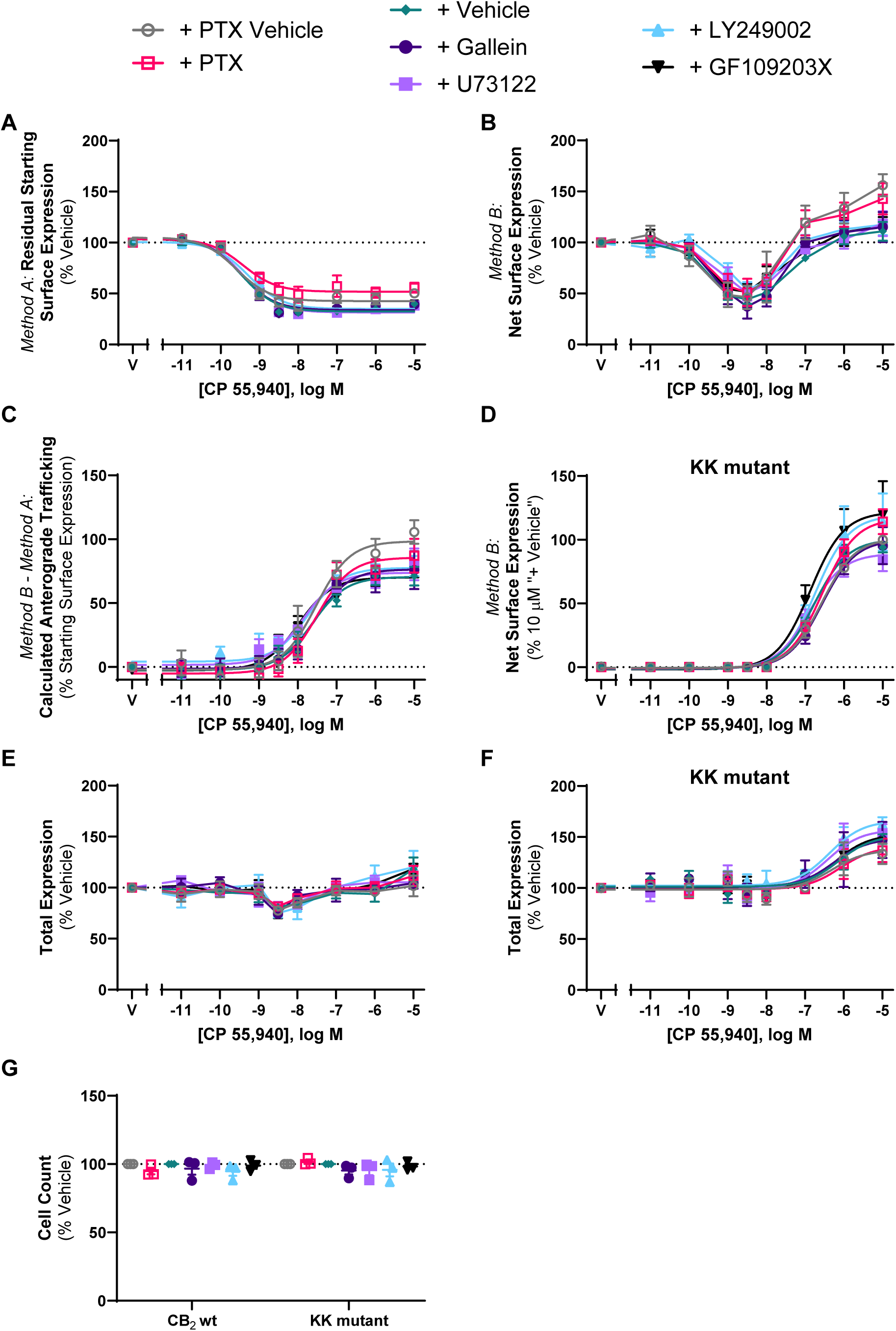
CB_2_ wt and KK mutant trafficking in response to CP 55,940 in the presence of various signalling inhibitors HEK cells stably expressing CB_2_ wt (A, B, C, E) or KK mutant (D, F) were treated with a CP 55,940 dilution series for 3 h, in the absence or presence of various signalling inhibitors, and were labelled to measure. (A) residual starting surface expression (*Method A*), (B, D) net surface expression (*Method B*), or (E, F) total receptor expression. (C) Calculated CB_2_ wt anterograde trafficking (*Method B* – *Method A*). (G) Average cell count from panels E and F. Data are presented as mean ± SEM from three independent experiments (except inhibitor vehicle in A-D, n=6).

**Table 3.** Trafficking summary data for stably expressed CB_2_ and KK mutant in response to CP 55,940, in the absence or presence of various inhibitors of signalling, trafficking, or protein synthesis. Parameters are derived from three independent experiments, unless otherwise stated.

|  | *Method A:*<br>Internalisation (3 hour) |  | *Method B – Method A:*<br>CP 55,940-Induced Anterograde Trafficking (3 hour) |  |  |  |
| --- | --- | --- | --- | --- | --- | --- |
|  | CB<sub>2</sub> wt |  | CB<sub>2</sub> wt |  | KK mutant |  |
|  | pEC<sub>50</sub><br>(± SEM) | E<sub>max</sub>, %<br>Vehicle<br>(± SEM) | pEC<sub>50</sub><br>(± SEM) | E<sub>max</sub>, % Starting<br>Surface<br>Expression,<br>Vehicle (± SEM) | pEC<sub>50</sub><br>(± SEM) | E<sub>max</sub><sup>#</sup><br>(± SEM) |
| **+ PTX Vehicle** | 9.60 (0.05) | -62.7 (4.9) | 7.56 (0.24) | 102.4 (8.6) | 6.59 (0.12) | 102.1 (2.4) |
| **+ PTX**<br>**(Gα<sub>i/o</sub> & Gβγ**<br>**inhibitor)** | 9.40 (0.02) | -52.3 (9.4) | 7.42 (0.07) | 90.1 (7.4) | 6.59 (0.08) | 117.3 (12.5) |
| **+ Vehicle\*** | 9.46 (0.05) | -71.6 (1.7) | 7.64 (0.13) | 73.8 (8.1) | 6.75 (0.10) | 102.9 (2.2) |
| **+ Gallein**<br>**(Gβγ inhibitor)** | 9.47 (0.05) | -71.1(2.8) | 7.58 (0.14) | 78.4 (13.0) | 6.57 (0.09) | 101.2 (13.9) |
| **+ U73122**<br>**(PLC inhibitor)** | 9.42 (0.02) | -72.5 (1.8) | 7.83 (0.12) | 72.6 (7.3) | 6.87 (0.10) | 90.8 (10.8) |
| **+ LY249002**<br>**(PI3K inhibitor)** | 9.43 (0.02) | -66.7 (1.5) | 7.74 (0.14) | 73.1 (4.0) | 6.72 (0.03) | 120.5 (23.7) |
| **+ GF109203X**<br>**(PKC inhibitor)** | 9.45 (0.05) | -71.1 (2.2) | 7.96 (0.15) | 73.2 (1.8) | 6.87 (0.13) | 124.3 (23.8) |
| **+ Vehicle<sup>^</sup>** | 9.48 (0.04) | -67.9 (3.9) | 7.30 (0.18) | 71.6 (14.3) | 6.60 (0.04) | 104.3 (2.1) |
| **+ BFA** | 9.38 (0.04) | -75.5 (3.1) | - | - | - | - |
| **+ Monensin** | 9.45 (0.02) | -75.7 (4.2) | 7.35 (0.18) | 36.4 (8.0) | - | 2.7 (1.0) |
| **+ CHX** | 9.56 (0.06) | -74.1 (1.1) | 7.20 (0.12) | 29.3 (11.4) | 6.18 (0.15) | 13.1 (3.0) |

<sup>#</sup> KK mutant CP 55,940-induced anterograde trafficking E<sub>max</sub> is normalised to the efficacy measured at 10 μM for the “+ Vehicle” conditions. \* All parameters derived from 6 independent experiments. <sup>^</sup> All parameters derived from 4 independent experiments.

### 3.5 CP 55,940-induced anterograde trafficking of CB_2_ is blocked by inhibitors of Golgi export, vesicle acidification, and protein synthesis

To further characterise CP 55,940-stimulated surface delivery of CB_2_, the effect of inhibitors of receptor trafficking and protein synthesis were investigated. Brefeldin A (BFA) is a macrocyclic lactone that blocks coat protein complex I (COPI) assembly at the Golgi by inhibiting ADP-ribosylation factor (ARF)1, thereby disrupting the Golgi and preventing protein secretion to the plasma membrane [51]. Monensin is an ionophore that facilitates the transmembrane exchange of sodium ions for protons, resulting in the neutralisation of acidic organelles, including the Golgi and some types of endosomes, and subsequently inhibiting receptor trafficking to the cell surface [52]. CHX inhibits protein synthesis at the translational level [53] and was included to determine whether new protein synthesis was required for this phenotype to occur.

These inhibitors did not influence CB_2_ internalisation in response to CP 55,940 (*p* > 0.05; Fig. 8A, Table 3). However, co-administration of these inhibitors, did influence net surface expression and the resulting extent of anterograde trafficking in response to CP 55,940 (Fig. 8B-D).

**Fig. 8.**
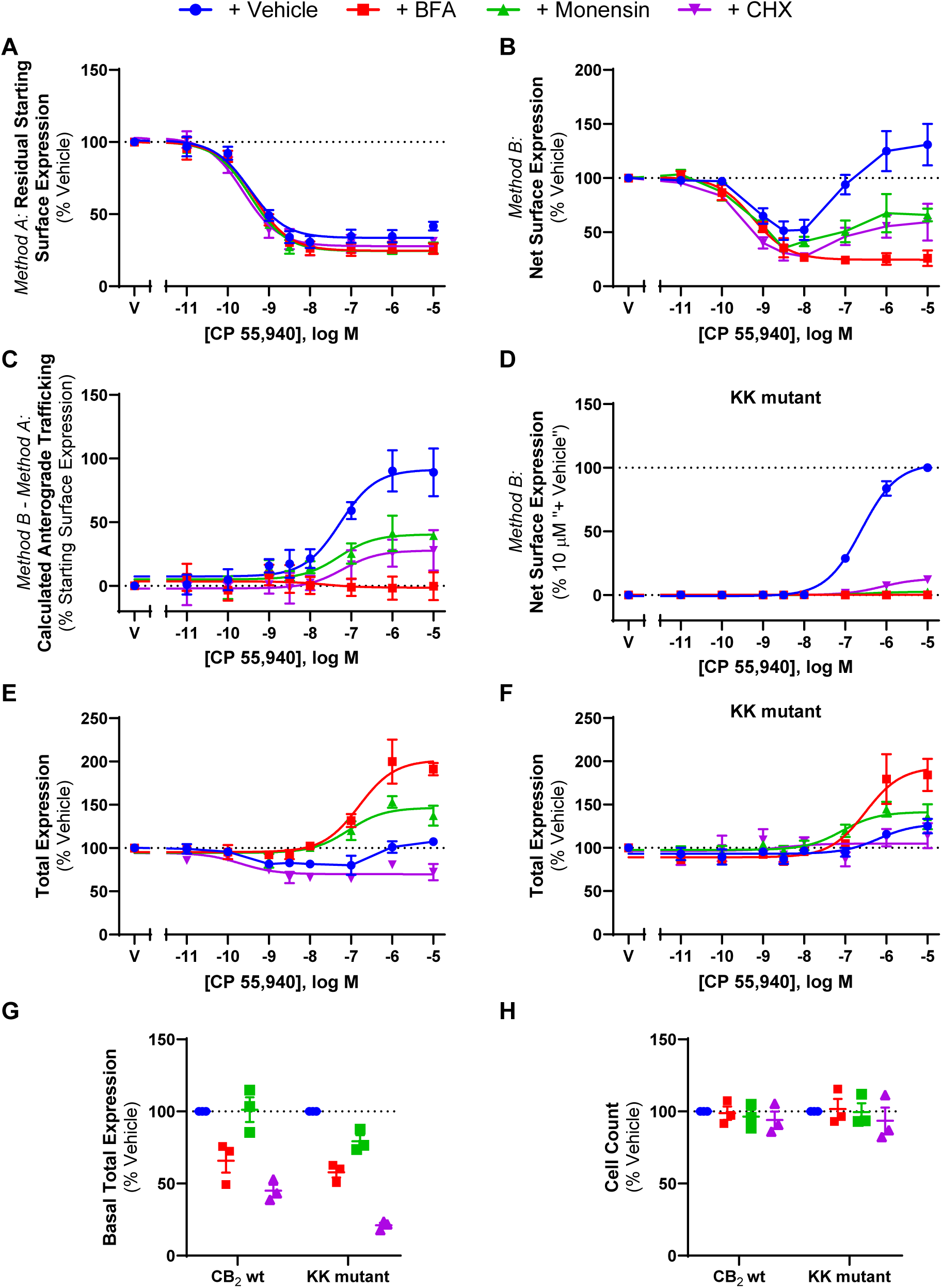
CB_2_ wt and KK mutant trafficking in response to CP 55,940 in the presence of various trafficking-related inhibitors HEK cells stably expressing CB_2_ wt (A, B, C, E) or KK mutant (D & F) were treated with a CP 55,940 dilution series for 3 hours, in the absence or presence of trafficking or protein synthesis inhibitors, and were labelled to measure. (A) residual starting surface expression (*Method A*), (B, D) net surface expression (*Method B*), or (E, F) total receptor expression. (C) Calculated CB_2_ wt anterograde trafficking (*Method B* – *Method A*). (G) Basal total expression, and (H) average cell count from ‘G’. Data are presented as mean ± SEM from three independent experiments.

Treatment with BFA reduced CB_2_ wt net surface expression in response to CP 55,940, producing a concentration-response curve in this labelling condition (Fig. 8B) with equivalent efficacy and potency to that when measuring internalisation alone (Fig. 8A). CP 55,940-induced anterograde trafficking was therefore found to be abolished when BFA was present (Fig. 8C). This was also apparent with the KK mutant where CP 55,940 did not stimulate CB_2_ wt surface delivery in the +BFA condition (Fig. 8D).

The presence of monensin similarly attenuated CB_2_ wt net surface expression in response to CP 55,940 (Fig. 8B), and CP 55,940-induced anterograde trafficking was partially blocked by this inhibitor, wherein a ∼50% reduction in efficacy relative to the vehicle-treated condition was observed (Fig. 8C, Table 3; *p =* 0.018). This was evident with the KK mutant also, but to an even greater degree, where CP 55,940-induced surface delivery was completely blocked (Fig. 8D, Table 3).

Inhibiting protein synthesis with CHX treatment also resulted in a significant decrease in net surface expression of CB_2_ wt following CP 55,940 stimulation, and partial blockade of the stimulated surface delivery phenotype, reducing its efficacy by ∼60% when compared to CP 55,940 plus inhibitor vehicle (Fig. 8B & C, Table 3; *p =* 0.026). Again, this was recapitulated with the KK mutant, where the efficacy of CP 55,940-induced CB_2_ wt anterograde trafficking was reduced to ∼15% of the CP 55,940 plus inhibitor vehicle response (Fig. 8D, Table 3).

A significant reduction in CB_2_ basal total expression was found in the presence of CHX (*p =* 0.001) and BFA (*p =* 0.010), but not monensin (Fig. 8G). For the KK mutant, all inhibitors produced a significant reduction in basal total expression relative to vehicle (*p =* 0.002 for BFA, *p =* 0.043 for monensin, *p <* 0.001 for CHX). Additionally, cell numbers were consistent between conditions, indicating that none of the inhibitors had a gross effect on cell health/viability (*p* > 0.05; Fig. 8H). When normalised to their respective CP 55,940 vehicles, CB_2_ wt total expression plus inhibitor vehicle was slightly reduced in response to intermediate concentrations of CP 55,940, but a lesser/no change was observed at higher concentrations (Fig. 8E). The presence of monensin resulted in an increase in CB_2_ wt total expression, reaching a maximal ∼30% above CP 55,940 plus vehicle, but this was not statistically significant (*p* > 0.05). Co-incubation with BFA significantly increased CB_2_ wt total expression at concentrations of CP 55,940 that would typically induce CB_2_ wt anterograde trafficking, reaching a maximum efficacy above CP 55,940 plus vehicle of ∼85% (*p =* 0.013). Conversely, when CHX was present, CP 55,940 decreased total CB_2_ wt expression (*p =* 0.036). KK mutant total expression data similarly showed a slight increase in total expression in response to CP 55,940 plus vehicle, and this was further increased in the presence of BFA, however this was not significant (*p* > 0.05). CHX and monensin similarly did not affect KK mutant total expression in response to CP 55,940 (*p* > 0.05).

### 3.6 CB_2_ anterograde trafficking is induced by cannabinoid agonists, antagonists, and inverse agonists

The data presented above suggests that CP 55,940-induced anterograde trafficking is not signalling-mediated, involves Golgi trafficking, requires new protein synthesis, and is specific to CB_2_. In combination with the lipophilic nature of CP 55,940 (and cannabinoids in general) and the CP 55,940-mediated changes observed in the western blots, the evidence so far is consistent with CP 55,940-stimulated anterograde trafficking being due to pharmacological chaperoning. As pharmacological chaperoning of GPCRs has previously been observed in response to both agonists and inverse agonists [54,55], we investigated whether other CB_2_ ligands, including inverse agonists, had the same effect as CP 55,940.

Trafficking assays were carried out in response to a panel of CB_2_ agonists using the aforementioned labelling methods (*Methods A* and *B*). The ligands chosen were structurally diverse and spanned various cannabinoid classes. Synthetic cannabinoids HU308 and WIN 55,212-2 all internalised CB_2_ wt when labelled *prior* to drug stimulation (*Method A*, Fig. 9A-B, Table 4). CB_2_ wt internalisation was not observed in response to the phytocannabinoid Δ^9^-THC (Fig. 9C), despite acting as a partial agonist in a cAMP assay (Supplementary Fig. 2). Endocannabinoids AEA and 2-AG both internalised CB_2_ wt when labelled via *Method A*, with AEA internalising to a lesser extent (Fig. 9D and E, Table 4). As these ligands possess a wide range of affinities for CB_2_, significant differences in internalisation EC_50s_ were expected, and this was accordingly observed (*p <* 0.001). In regard to efficacy, significant differences were observed between AEA vs. 2-AG and CP 55,940 (*p =* 0.016 and *p =* 0.037, respectively), and WIN 55,212-2 vs. 2-AG and CP 55,940 (*p =* 0.007 and *p =* 0.016). As Δ^9^-THC did not internalise CB_2_ wt, it was not included in this statistical analysis.

**Fig. 9.**
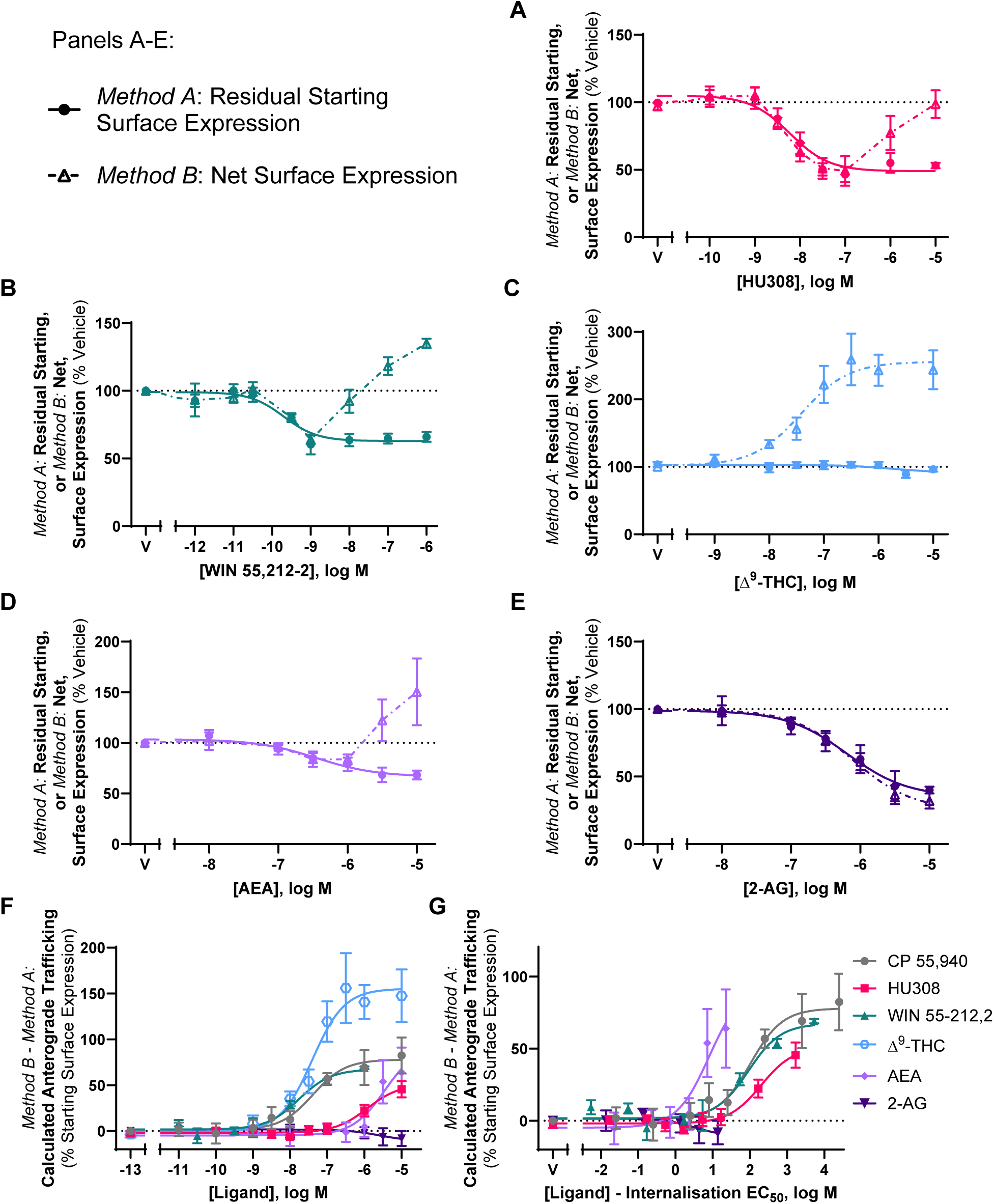
CB_2_ wt trafficking in response to a panel of CB_2_ agonists Cells stably expressing CB_2_ wt were treated with dilution series of various CB_2_ agonists for 3 hours and labelled to measure. (A-E) residual starting surface expression (*Method A*) or net surface expression (*Method B*). (F) Calculated ligand-induced CB_2_ wt anterograde trafficking (*Method B* – *Method A* from panels A-E; CP 55,940 re-plotted from Fig. 1D). (G) Calculated ligand-induced anterograde trafficking from panel F, normalised on the x-axis to each ligand’s respective internalisation EC_50,_ wherein the internalisation logEC_50_ was subtracted from the treatment concentrations. Data are presented as mean ± SEM from three independent experiments (except AEA, n=4).

**Table 4.** Trafficking summary data for stably expressed CB_2_ and KK mutant in response to various CB_2_ ligands. Parameters are derived from three independent experiments, unless otherwise stated.

|  | *Method A: Internalisation (3 hour)* |  | *Method B – Method A:*<br>*Calculated Anterograde Trafficking (3 hour)* |  |  | *Method B: Net Surface*<br>*Expression (3 hour)* |  |
| --- | --- | --- | --- | --- | --- | --- | --- |
|  | CB<sub>2</sub> wt |  | CB<sub>2</sub> wt |  |  | KK mutant |  |
|  | pEC<sub>50</sub> (± SEM) | E<sub>max</sub> (± SEM)<sup>a</sup> | pEC<sub>50</sub> (± SEM) | Internalisation-Adjusted EC<sub>50</sub><br>(± SEM)<sup>b</sup> | E<sub>max</sub> (± SEM)<sup>c</sup> | pEC<sub>50</sub><br>(± SEM) | E<sub>max</sub><br>(± SEM)<sup>d</sup> |
| **CP 55,940<sup>°</sup>** | 9.40 (0.09) | -64.4 (2.4) | 7.35 (0.17) | 2.05 (0.25) | +76.6 (10.0) | N.D.<sup>+</sup> | N.D.<sup>+</sup> |
| **HU308** | 8.23 (0.10) | -56.3 (3.1) | 5.91 (0.19) | 2.32 (0.29) | +53.7 (7.0) | N.D. | N.D. |
| **WIN 55,212-2** | 9.73 (0.12) | -37.3 (1.6)* | 7.76 (0.16) | 1.97 (0.08) | +65.7 (4.6) | N.D. | N.D. |
| **Δ<sup>9</sup>-THC** | Not measurable | -3.7 (3.6)<sup>^</sup> | 7.43 (0.16) | Not measurable | +158.5 (32.5)** | N.D. | N.D. |
| **AEA** | 6.35 (0.16) | -42.1 (6.0)* | Not measurable | Not measurable | +63.9 (27.1)<sup>#</sup> | N.D. | N.D. |
| **2-AG** | 6.13 (0.20) | -67.6 (4.1) | Not measurable | Not measurable | -8.2 (8.2)<sup>^</sup> | N.D. | N.D. |
| **AM630** | 8.17 (0.14) | +35.1 (6.0) | 7.36 (0.06) | 0.81 (0.18) | +238.9 (10.8) | 6.29 (0.12) | 209.3 (20.6) |
| **SR144528** | 8.15 (0.19) | +25.9 (1.2) | 7.82 (0.20) | 0.33 (0.39) | +229.5 (2.8) | 6.37 (0.15) | 178.4 (10.7) |

N.D. = Not determined. <sup>a</sup> Internalisation E<sub>max</sub> represented as reduction (–) or increase (+) in surface expression, as a percentage of vehicle-treated control.
<sup>b</sup> Internalisation-adjusted anterograde trafficking EC<sub>50</sub> calculated for each ligand by subtracting the EC<sub>50</sub> for internalisation from the anterograde trafficking EC<sub>50</sub>.
<sup>c</sup> Anterograde trafficking was calculated by subtracting internalisation (*Method A*, normalised to vehicle-treated control) from net surface expression (*Method B*, normalised to vehicle-treated control). E<sub>max</sub> is therefore represented as reduction (–) or increase (+), as a percentage of starting surface expression. <sup>d</sup> KK mutant net surface expression E<sub>max</sub> is normalised to that induced by 10 μM CP 55,940 at the same timepoint.
<sup>°</sup> CP 55,940 data for CB<sub>2</sub> wt is re-tabulated from Table 1 (adding new calculation for internalisation-adjusted EC<sub>50</sub>). <sup>#</sup> A fully defined concentration-response curve could not be obtained, therefore efficacy at the highest concentration tested (i.e., 10 μM) is noted instead of E<sub>max</sub>.
<sup>^</sup> A concentration-response curve could not be fitted, therefore efficacy at the highest concentration tested (i.e., 10 μM) is noted instead of E<sub>max</sub>.
<sup>+</sup> Not determined in this experiment set. See Table 2 for data. \* *p* < 0.05 vs. CP 55,940 and 2-AG. \*\* *p* < 0.05 vs. HU308, WIN 55,212-2, 2-AG and AEA.

When labelled *following* drug stimulation (*Method B*), the synthetic cannabinoids induced a response similar to CP 55,940, whereby CB_2_ wt was internalised at lower agonist concentrations, but at greater concentrations net surface expression was unchanged or increased (Fig. 9A-B). Despite not internalising CB_2_ wt, Δ^9^-THC increased net surface expression in a concentration-dependent manner, reaching a maximal efficacy of ∼160% above vehicle (Fig. 9C). AEA produced a similar phenotype to the synthetic cannabinoids when labelled via *Method B* (Fig. 9D). Conversely, 2-AG internalised CB_2_ in a concentration-dependent manner, producing concentration-response curves with equivalent efficacy (40.1 [± 2.5] vs. 31.9 [± 5.69]) and potency (6.13 [± 0.20] vs. 6.08 [± 0.17]) in both labelling conditions (Fig. 9E). A possible explanation for 2-AG not inducing CB_2_ anterograde trafficking is that it may be degraded and therefore unable to interact with intracellular CB_2_. To address this, trafficking assays with 2-AG were carried out in the presence and absence of JZL184, an inhibitor of monoacylglycerol lipase (MAGL, the metabolising enzyme that 2-AG is primarily degraded by). CB_2_ wt internalisation in response to 2-AG was unaffected by JZL184 (*p* > 0.05), and no anterograde trafficking was detected in the presence of JZL184 (Supplementary Fig. 2).

For the panel of CB_2_ agonists, ligand-induced anterograde trafficking was calculated by subtracting the surface expression detected via labelling *Method A* from the surface expression detected via labelling *Method B*. As seen in Fig. 9F and Table 4, the above ligands induced anterograde trafficking of CB_2_ wt with varying potencies, which is unsurprising given the range of internalisation potencies observed. While most of the ligands screened induced anterograde trafficking of CB_2_ wt to similar extents as CP 55,940 (∼55-80% of starting surface expression), Δ^9^THC stimulation produced a greater efficacy of ∼160%, whereas 2-AG did not induce this phenotype at all. To compare the propensity of each ligand to induce CB_2_ anterograde trafficking, these concentration-response curves were re-plotted on the x-axis such that each ligand’s own internalisation EC_50_ was subtracted from each measured concentration (Fig. 9G). If the resulting internalisation-adjusted EC_50_ for a compound were 0, this would indicate its propensity to induce anterograde trafficking was equivalent to that of internalisation. If the internalisation-adjusted EC_50_ were > 0, the propensity of the compound to induce anterograde trafficking would be lesser than that for internalisation. This calculation therefore allows comparison between ligands independent of internalisation potency. CP 55,940 was also included in this analysis. As Δ^9^-THC did not internalise hCB_2_ wt and the anterograde trafficking EC_50_ for AEA could not be measured reliably, these ligands were not included in the statistical analysis (Table 4) though, when plotted, AEA qualitatively trended toward having the highest potency for anterograde trafficking relative to internalisation EC_50_ of the ligands tested (Fig. 9G). HU308 qualitatively trended toward having lower relative potency for hCB_2_ wt anterograde trafficking than CP 55,940 and WIN 55,212-2, but no statistically significant difference between these ligands was found (*p* = 0.55).

The above trafficking assays were also carried out in response to CB_2_-selective inverse agonists, AM630 and SR144528. When CB_2_ wt was labelled *prior* to inverse agonist simulation, both inverse agonists produced a concentration-dependent increase in staining relative to vehicle (indicative of inhibiting constitutive internalisation) with equivalent potencies and similar efficacies (Fig. 10A and B, Table 4). When CB_2_ wt was labelled *following* inverse agonist stimulation, concentration-dependent increases in surface expression were again measured in response to AM630 and SR144528, but to much greater extents (Fig. 10A and B, Table 4). The efficacies produced here are also considerably larger than were produced by agonists, as seen when compared to CP 55,940 (Fig. 10A and B, grey line).

**Fig. 10.**
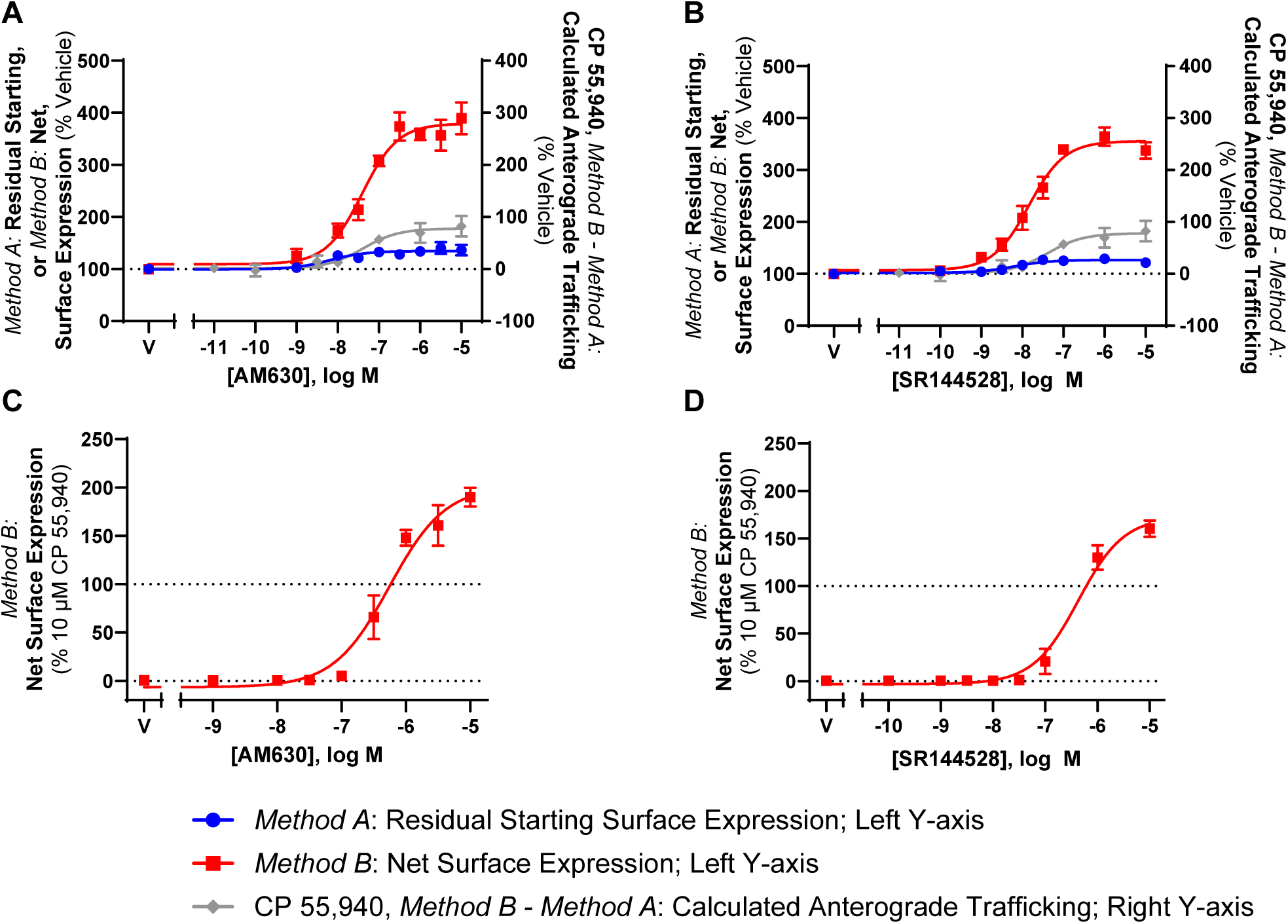
CB_2_ wt and KK mutant trafficking in response to CB_2_ inverse agonists Left Y-axis: Cells stably expressing. (A,B) CB_2_ wt or (C,D) KK mutant were treated with dilution series of CB_2_-selective inverse agonists, AM630 and SR 144528, for 3 hours, and were labelled to measure residual starting surface expression (*Method A*) or net surface expression (*Method B*). Right Y-axis: In A-B, calculated CP 55,940-induced anterograde trafficking (*Method B – Method B*) is shown for reference. Data are presented as mean ± SEM from three independent experiments.

To confirm that the inverse agonist-induced increases in CB_2_ wt surface expression observed in Fig. 10A and B were not only a result of the stabilisation of existing surface receptors and/or the prevention of constitutive degradation, trafficking assays in AM630 and SR144528 were carried out with the KK mutant. Surface receptor expression was labelled *following* drug stimulation, and 1 µM CP 55,940 was included for reference. Indeed, surface expression of the KK mutant was induced in response to both inverse agonists, reaching a maximal efficacy ∼110% greater than that induced by CP 55,940 for AM630, and ∼80% greater for SR144528, and confirming that these ligands are inducing CB_2_ anterograde trafficking (Fig. 10C and D, Table 4). Inverse agonist-induced anterograde trafficking of CB_2_ was ∼1 log unit more potent at CB_2_ wt vs. the KK mutant for AM630, and ∼1.5 log units more potent at CB_2_ wt for SR144528. Trafficking summary parameters for all ligands in this section are presented in Table 4.

## 4 Discussion

Herein, we have presented evidence that CB_2_ surface expression can be stimulated by application of lipophilic CB_2_ agonists and inverse-agonists that likely act as pharmacological chaperones. As well, we report that a di-lysine (KK) motif in the C-terminal tail is required for constitutive, but not chaperoned, delivery of CB_2_ to the cell surface.

Initially studying CP 55,940 as a prototypic CB_2_ agonist, pharmacological chaperoning was demonstrated to be particular to CB_2_ as this was not observed for CB_1_ or B_2_AR. These experiments also confirmed that CP 55,940-induced CB_2_ surface delivery was not due to modulation of the CMV promoter, nor an artefact of the labelling methods/antibodies employed. Furthermore, in the presence of BFA to block release of CB_2_ from the secretory pathway, a treatment used previously to attenuate pharmacological chaperoning of other GPCRs [56,57], equivalent internalisation responses were measured whether antibody was applied before (*Method A*) or after (*Method B*) drug stimulation. This again validated that CB_2_ internalisation observed utilising *Method A* is not an artefact of live cell antibody labelling. The potential for CB_2_-mediated signalling to contribute CP 55,940-induced surface delivery was ruled out by the lack of effect of a panel of signalling pathway inhibitors.

Providing further evidence for a pharmacological chaperone effect, at the same time as BFA blocked agonist-induced CB_2_ cell surface delivery, it considerably enhanced the agonist-induced increase in total expression. This indicates that CP 55,940 has a positive effect on CB_2_ expression stability prior to exit from the secretory pathway, likely preventing degradation by quality control machinery [17]. Furthermore, these observations imply that when BFA is *not* present, increases in surface and total expression in response to CP 55,940 arise primarily from increased receptor stability in the secretory pathway, rather than post-delivery stabilisation at the plasma membrane or prevention of post-endocytic degradation.

Interestingly, agonist-stimulated CB_2_ surface delivery was partially blocked by the protein synthesis inhibitor CHX. This implies that the majority of chaperoned CB_2_ is not derived from the pre-existing intracellular pool. Instead, chaperoning ligand needs to be present at a certain stage of CB_2_ synthesis or maturation to be able to promote delivery to the cell surface. This is also consistent with time course data, where an initial lag time of at least one hour before a significant increase in CB_2_ anterograde trafficking in response to CP 55,940 is observed. Although CHX (as a broad spectrum protein synthesis inhibitor) could have blocked anterograde receptor trafficking via reducing expression of chaperone proteins, this seems unlikely as the half times for relevant chaperone proteins are reportedly in the scale of multiple hours [58,59]. These conclusions from experiments with CHX and BFA parallel earlier findings for the delta opioid receptor (DOR), wherein a large proportion of DOR was found not to be delivered to the cell surface and was instead proteasomally degraded, but application of cell permeable antagonist could prevent this degradation and instead promote cell surface delivery [55].

Early reports of GPCR pharmacological chaperoning focused on disease-causing mutations with low basal cell surface expression that could be rescued by application of lipophilic receptor ligands [17]. While we are not aware of any naturally occurring or disease-associated SNPs or mutations that markedly alter CB_2_ surface expression [25], we tested whether motifs in the C-terminal tail with strong potential to be implicated in membrane protein trafficking are relevant for CB_2_. Despite the well-established role of conserved cytosolic/C-terminal di-acidic motifs ([D/E]*X*[D/E]) interacting with coat protein complex II (COPII) machinery and mediating ER export of several non-GPCR transmembrane proteins (e.g. vesicular stomatitis virus glycoprotein [VSV-G] [43,44], cystic fibrosis transmembrane conductance regulator [CFTR] [45]), and the angiotensin II type 2 receptor (AT_2_R, [41]), ETE339-41ATA had no measurable effect on CB_2_ trafficking. Truncation of the last 12 C-terminal residues that includes a phosphorylation site (S352) [5], a cluster of aspartic acid residues which may act as phospho-mimics [46], and a type 2 PDZ-binding motif [47], also had surface expression and ligand-stimulated trafficking that were equivalent to CB_2_ wt. Interestingly, total expression of the Δ12 mutation was enhanced, implying increased intracellular expression.

Conversely, the KK318-319EE mutation inhibited all basal surface expression, despite having similar total expression to wild-type CB_2_. Classically, a di-lysine ‘KK’ motif on transmembrane proteins serves as a retrieval motif, interacting with coat protein complex I (COPI) to facilitate retrograde transport from the Golgi to the ER [60,61]. However, this appears to have a context-dependent role in GPCRs, acting as an ER *retention* motif in metabotropic glutamate receptor 1b (mGluR1b) [39], but shown to be important for surface expression of angiotensin II type 1 receptor (AT_1_R; [42]) and bitter taste receptor 4 [40], consistent with our observations here for CB_2_. Antibody “feeding” experiments clarified that the lack of CB_2_ surface expression was due to lack of receptor delivery to the cell surface, rather than rapid internalisation. Interestingly, KK318-9 is conserved between human and rodent CB_2_, despite considerable heterogeneity between the C-terminal tail sequences [62]. Only limited data is available on the specific role of the KK motif in GPCR trafficking, but this has been suggested to interact with tubulin, which might explain prevention of basal surface trafficking [42]. However, in strong support of the pharmacological chaperoning hypothesis, CP 55,940 concentration-dependently “rescued” cell surface expression of the CB_2_ KK318-9EE mutant. This implies that either pharmacological chaperoning by CB_2_ ligands can divert CB_2_ to be delivered to the cell surface via a different mechanism from basal wt receptor delivery that is not dependent on an intact KK motif, or that the KK318-9EE mutation introduces a ‘gain of function’ retention of CB_2_ in the secretory pathway that is reversed by chaperoning to enable CB_2_ delivery to the cell surface.

A hallmark of pharmacological chaperoning in other GPCRs is a shift in receptor maturation state from immature to mature species with chaperoning, as observed via western blotting (e.g. [21,55]). Here, we find that basally a considerable population of CB_2_ resolves in a ∼67 kDa band, with additional species at a range of larger sizes (perhaps complexed with chaperone proteins) and that the KK mutant has a larger proportion in these species in comparison with wt. In response to CP 55,940 stimulation, the relative proportion of these large complexes decreased and an increase in ∼40 to ∼55 kDa species was observed for both CB_2_ wt and KK mutant, which likely corresponds to the receptor being released from larger complexes and/or undergoing post-translational modifications then being delivered to the cell surface. Interestingly, the proportions of ∼35 and 40 kDa (likely immature) species were similar between CB_2_ wt and KK mutant and unaffected by CP 55,940 stimulation, perhaps representing the intracellular pool.

Importantly, a panel of diverse CB_2_ ligands – both agonists and inverse-agonists – were found to act as CB_2_ pharmacological chaperones. Agonists HU308, WIN 55,212-2, AEA and 2-AG all internalised CB_2_ wt, with partial efficacy for WIN 55,212-2 and AEA being consistent with literature showing partial agonism for these ligands in β-arrestin 2 recruitment [26,63]. Δ^9^-THC did not induce CB_2_ internalisation, which was consistent with a prior-reported lack of β-arrestin recruitment [63,64]. Despite this, Δ^9^-THC did engage with CB_2_ as it acted as a partial agonist for inhibition of forskolin-stimulated cAMP production and stimulated cell surface expression to a greater overall extent than CP 55,940 and the other agonists tested. The effect on cell surface expression raises the possibility that a contributing mechanism to the immunomodulatory effects of cannabis may be via enhancing CB_2_ expression, after which endocannabinoids might induce greater effects. When the potency of ligand-induced CB_2_ anterograde trafficking was assessed relative to each agonist’s internalisation potency, no significant differences were observed between CP 55,940, HU308, and WIN 55,212-2. This suggests that these ligands all have a similar propensity to induce CB_2_ anterograde trafficking relative to their ability to induce internalisation, with potencies ∼75-to 200-fold lower than that for internalisation – this shift perhaps being related to the lag time prior to onset of stimulated anterograde trafficking. While a reliable EC_50_ could not be determined for AEA (as no plateau was observed), this qualitatively appeared to be the most potent of all the ligands screened for inducing anterograde trafficking. Despite acting as a full agonist for internalisation, 2-AG did not stimulate anterograde trafficking within the range of concentrations tested after 3 hours of stimulation. We verified this was unlikely to be due to enzymatic degradation of 2-AG during the assay via inclusion of a MAGL inhibitor. However, of the ligands tested 2-AG had the lowest potency for inducing internalisation and, based on relative ligand-stimulated anterograde trafficking potencies observed for most ligands (other than AEA), the highest concentration applied might not have been sufficient to produce chaperoning even if 2-AG has the potential to act in a similar way as the other agonists.

Inverse agonists AM630 and SR144528 increased surface expression of hCB_2_ wt, which was consistent with previous studies [5,6]. These also induced CB_2_ KK mutant surface expression, indicating that the increase in wild-type CB_2_ surface expression was likely a result of both inhibition of constitutive internalisation and stimulated anterograde trafficking. Lack of induction of endocytosis might explain the overall greater efficacy of inverse agonists than agonists for inducing anterograde trafficking, assuming that agonists may induce a degree of endocytosis subsequent to receptor chaperoning to the cell surface. Alternatively, or potentially in addition, inverse agonist binding might stabilise a receptor conformation that is more amenable to chaperoning than agonists. The inverse agonist findings are also further evidence that the stimulated anterograde trafficking phenotype is not induced by agonist signalling and likely results from pharmacological chaperoning.

A limitation of our study is that experiments were performed in a heterologous expression system. It will be important to verify the ligand-induced anterograde trafficking phenotype in cells endogenously expressing CB_2_, and investigate both the acute and long-term functional consequences. This may be challenging due to widely-reported poor reliability of anti-CB_2_ antibodies [65–67] and because alternative ligand-based methods, such as radioligand binding or fluorescent ligand binding, are unlikely to be practical for validating this trafficking phenotype due to ligand competition issues. Furthermore, while other studies investigating pharmacological chaperoning of GPCRs have included the use of membrane-impermeant ligands as negative controls (e.g. [21,55]), there are currently no confirmed membrane-impermeant CB_2_ ligands available to test, limiting our ability to delineate surface versus intracellular functional consequences. There are, however, *less* lipophilic CB_2_ ligands available that would be interesting to investigate [3,37,68], and the KK318-9EE CB_2_ mutant will likely serve as a useful tool in future studies to delineate functions of different CB_2_ receptor pools.

The findings of this study indicate that basal CB_2_ surface delivery is reliant on a C-terminal KK motif, and ligand-induced anterograde trafficking of CB_2_ occurs via an independent pathway that is not downstream of G protein activation, but is likely the result of CB_2_ ligands acting as pharmacological chaperones on newly synthesised CB_2_. While a range of both agonists and inverse agonists induced this phenotype, it is notable that two specific physiologically/clinically relevant ligands, 2-AG and Δ^9^-THC, were unique. The interplay between the effects of endogenous and exogenous ligands on CB_2_ subcellular distribution and consequent signalling from different compartments may have critical impacts on both short– and long-term CB_2_-mediated responses. Meanwhile, the potential for CB_2_ ligands to interact with intracellular receptors is important to consider in the context of medicinal chemistry efforts where optimisation of physicochemical properties may alter ligand membrane permeability. Further characterisation such as identifying chaperone/adaptor proteins involved may also provide novel therapeutic targets for modulating CB_2_ subcellular localisation and, therefore, function.

## Statements and Declarations

### Author Contributions

Conceptualisation: NG, MG, CO; Investigation: CO, BW, NG, KW; Methodology: NG, CO; Visualization: CO, KW, NG; Supervision: NG, MG; Funding acquisition: NG; Project administration: NG; Writing – original draft: CO, NG; Writing – review & editing: All authors.

### Data Availability

The data that support the findings of this study are available from the corresponding author upon reasonable request.

### Ethics approval

Not applicable to this study

### Consent to participate

Not applicable to this study

### Consent to publish

Not applicable to this study.

### Author contributions

Conceptualisation: NG, MG, CO; Investigation: CO, BW, NG, KW; Methodology: NG, CO; Visualization: CO, KW, NG; Supervision: NG, MG; Funding acquisition: NG; Project administration: NG; Writing – original draft: CO, NG; Writing – review & editing: All authors.

## Acknowledgements

This research was funded by a Marsden Fast-Start grant from the Royal Society Te Apārangi New Zealand (UOA1507). CO was supported by a University of Auckland Doctoral Scholarship. NG was supported by Fellowships from the Auckland Medical Research Foundation (1313001) and the Health Research Council of New Zealand (20-006).

