## Supplementary Figures for "Insights into Cannabinoid Receptor 2 (CB2) anterograde trafficking and pharmacological chaperoning"

**Supplementary Fig. 1**

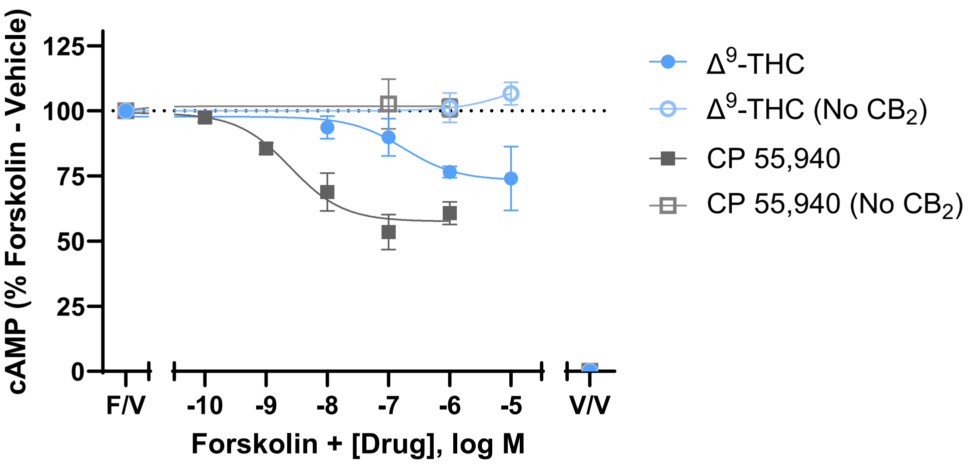


**Supplementary Fig. 1 CB$_2$ wt cAMP response to Δ$^9$-THC and CP 55,940**

Cells stably expressing CB$_2$ wt (or untransfected cells, "No CB$_2$") were treated with forksolin (5 μM) and a dilution series of Δ$^9$-THC or CP 55,940. Cyclic AMP (cAMP) responses were measured as the mean BRET ratio from the CAMYEL biosensor during a 10 minute stimulation, then normalised to forskolin with vehicle (100%; "F/V") and vehicle-only control (0%, "V/V"). Data are presented as mean ± SEM from three independent experiments.

### Supplementary Fig. 2

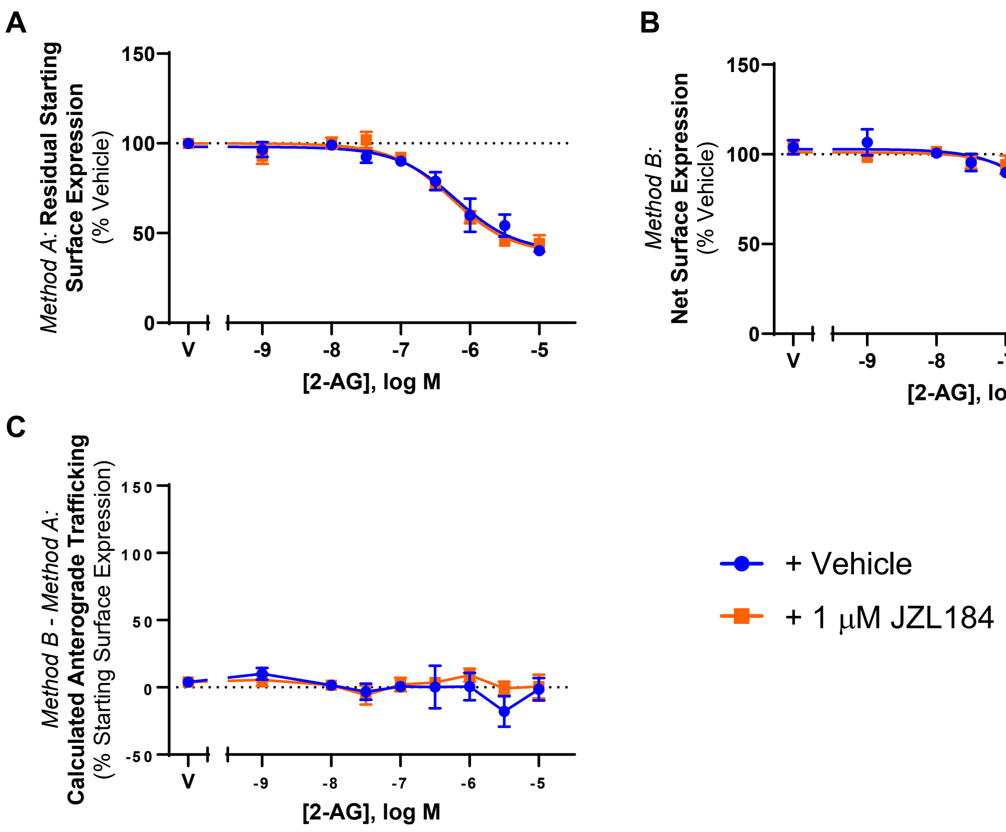


**Supplementary Fig. 2 $CB_2$ wt trafficking in response to 2-AG +/- MAGL inhibitor, JZL184**

Cells stably expressing $CB_2$ wt were treated with a dilution series of 2-AG +/- 1 μM JZL184 for 3 hours, and were labelled to measure (A) residual starting surface expression (*Method A*) or (B) net surface expression (*Method B*). (C) Calculated anterograde trafficking (*Method B – Method A*). Data are presented as mean ± SEM from three independent experiments.
